# Regulation of neuropathic pain by microglial Orai1 channels

**DOI:** 10.1101/2022.09.02.506250

**Authors:** Shogo Tsujikawa, Kaitlyn E DeMeulenaere, Marivi V Centeno, Shahrzad Ghazisaeidi, Megan E. Martin, Martinna R. Tapies, Mohammad M Maneshi, Megumi Yamashita, Kenneth A Stauderman, Apkar V Apkarian, Michael W Salter, Murali Prakriya

## Abstract

Microglia are important mediators of neuroinflammation that underlies neuropathic pain. However, the molecular checkpoints controlling microglial reactivity are not well-understood. We investigated the role of Orai1 channels for microglia-mediated neuroinflammation following nerve injury and find that deletion of Orai1 in microglia attenuates Ca^2+^ signaling and the production of inflammatory cytokines by proalgesic agonists. Conditional deletion of Orai1 attenuated microglia proliferation in the dorsal horn, spinal cytokines levels, and potentiation of excitatory neurotransmission following peripheral nerve injury. These cellular effects were accompanied by mitigation of pain hyperalgesia in Orai1 knockout mice. A small-molecule Orai1 inhibitor, CM4620, similarly mitigated allodynia in male mice. Surprisingly, these protective effects were not seen in female mice, revealing striking sexual dimorphism in Orai1 regulation of microglial reactivity and hyperalgesia. These findings indicate that Orai1 channels are key regulators of the sexually dimorphic role of microglia for the neuroinflammation that underlies neuropathic pain.

## INTRODUCTION

Microglia are the tissue-resident macrophage cells of the nervous system comprising about 5-10% of the cells in the central nervous system (*1*). Widely considered to be the sentinels of the brain, microglia in the healthy brain survey their environment for changes in synaptic activity, tissue damage, and infections via their highly ramified processes. Following brain damage or infection, however, microglia are activated rapidly via a process termed microgliosis to mediate several additional major roles (*2*). First, they promote neuronal wellbeing by removing debris and dying cells via phagocytosis and efferocytosis (*3*). Second, they defend against infections and tissue damage by secreting pro-inflammatory and pro-resolving mediators. Although these dueling functions are well balanced in the healthy brain, under certain pathological conditions, the failure of checkpoints in inflammatory signaling cascades can transform microglia into a disease-associated form, causing persistent, damaging neuroinflammation (*3*-*6*). In this study, we examine a potential role for store-operated Orai1 channels in regulating the inflammatory functions of activated microglia in the context of neuropathic pain.

Microglial reactivity and activation following nerve injury is driven by a variety of membrane receptors, but purinergic receptors including the P2X and P2Y families (*4*, *7*-*12*) have received especially strong scrutiny in microglial activation that follows nerve injury. Prevailing views indicate that stimulation of these receptors by ATP turns on the p38-mitogen activated protein (MAP) kinase and NF-κb pathways in microglia, triggering synthesis and release of a variety of pro-inflammatory agents including IL-6, TNF⍺, and IL-1β, which are thought to potentiate excitatory synaptic transmission in the dorsal spinal horn (*13*-*15*) to remodel spinal circuits involved in neuropathic pain (*5*, *12*, *16*-*18*). In line with this evidence, inhibiting pro-inflammatory cytokines with specific inhibitors (*19*-*21*), broadly targeting microglial activation with tetracycline antibiotics such as minocycline, or destroying microglia with targeted immunotoxins all reduce allodynia and other endpoints of neuropathic pain (*22*-*24*). Puzzlingly, however, microglial contributions to neuropathic pain are highly sex-dependent (*23*, *25*, *26*). Attenuation of hypersensitivity and allodynia by broad-spectrum microglial inhibitors is reported to occur in male mice but not female mice (*23*, *25*-*29*). Whether this gender difference is due to sex-dependent variations in the spinal neuronal responses to nerve injury, spinal cytokine levels, or cell autonomous differences in microglia is widely debated and not well understood (*28*, *29*). More generally, despite the explosion of information about neuropathic pain in recent years, no breakthrough treatments have emerged and even the mechanisms of the pathogenesis itself are not fully established (*30*-*32*). The slow progress probably reflects a combination of factors including inadequate genetic tools for isolating specific microglial signaling pathways and the multifactorial nature of neuropathic pain with interactions between multiple types of cells, cytokines, intrinsic abnormalities, and environmental factors.

As electrically non-excitable cells, microglial excitability is primarily regulated by cellular Ca^2+^ elevations evoked by cell surface receptors (*33*, *34*). These Ca^2+^ signals stimulate numerous microglial functions including phagocytosis, calcineurin/NFAT-signaling, and TLR-4 mediated NF-kB activation (*33*, *35*, *36*). Ca^2+^ regulation of these processes is particularly well suited for its potential to be stimulated rapidly and modulated by other signaling pathways. Yet, our understanding the pathways mediating microglial Ca^2+^ signaling and their contributions to microglial reactivity and neuroinflammation remains poorly understood. There is evidence for Ca^2+^ release from intracellular stores (*33*). However, because release of Ca^2+^ from intracellular stores is typically coupled to activation of store-operated Ca^2+^ entry (SOCE) (*37*) and previous studies were not designed to discriminate between SOCE and store release, the specific physiological contributions of SOCE for microglial reactivity and neuroinflammation has not been critically evaluated in past work.

In most animal cells, SOCE serves as a central mechanism for mobilizing [Ca^2+^]_i_ elevations and is activated by the engagement of G-protein or tyrosine kinase receptors (*37*) stimulate Ca^2+^ release from endoplasmic reticulum (ER) stores. The ensuing depletion of ER stores stimulates the ER Ca^2+^ sensors, the STIM1/2, which in turn activate Ca^2+^ release-activated Ca^2+^ (CRAC) channels formed by the Orai proteins (*37*). Orai1, the best studied member of this family, is exquisitely Ca^2+^ selective, making it ideally suited for generating oscillatory and long-lasting Ca^2+^ signals needed for transcriptional and enzymatic cascades (*37*, *38*). In the nervous system, Orai1 is implicated in a growing list of functions including gliotransmitter release, synaptic plasticity, phagocytosis, chemotactic migration, and activation of NF-κB and MAPK (*39*-*43*). Recently, a whole-animal Orai1 knockout was shown to be resistant to carrageenan- and formalin induced pain, which was attributed to changes in neuronal excitability, especially in the dorsal horn neurons of the spinal cord (*44*). However, the contribution of microglial Orai1 channels for neuroinflammation that underlies neuropathic pain is unknown.

These findings led us to consider several questions. Can manipulation of microglial activation by blocking SOCE affect microglial inflammation following nerve injury. How does this affect excitatory synaptic transmission in the spinal cord? And what are the implications for neuropathic pain? In this study, we addressed these questions using a microglial-specific Orai1 knockout mouse and biochemical, electrophysiological, and behavioral assays. We find that Orai1 channels in microglia are critical for the synthesis and release of inflammatory mediators and play a central role in the potentiation of synaptic transmission in the spinal cord following nerve injury and the mechanical sensitivity that is the hallmark of neuropathic pain.

## RESULTS

### Orai1 is essential for SOCE in microglia

To address the role of microglial Orai1 channels in regulating microglial Ca^2+^ signaling, neuroinflammation, and neuropathic pain following peripheral nerve injury, we generated a microglial specific Orai1 knockout (KO) mouse line by crossing Orai1*^fl/fl^* mice with the inducible Cre line, Cx3CR1-Cre/ERT2 (*45*) for selective deletion of Orai1 in microglia. Homozygous Orai1*^fl/fl^ ^Cx3CR1-Cre/ERT2^* mice arising from this cross were born at mendelian ratios and did not differ from WT mice in terms of weight, litter size, or gross mobility. To knock out Orai1 expression, we used two approaches. *(i)* For the *in vitro* mechanistic studies, we isolated primary spinal cord microglia from Orai1*^fl/fl^ ^Cx3CR1-Cre/ERT2^* mice or wildtype controls from P0-P2 stage, cultured them for 10 days, and added tamoxifen or vehicle to the culture medium for Orai1 deletion (see Methods). RT-qPCR analysis indicated that after tamoxifen treatment, microglia from Orai1*^fl/fl Cx3CR1-Cre/ERT2^* mice showed marked loss of Orai1 mRNA in cultured microglia **(Fig. S1A,B)** indicating that Cre recombination is highly successful in deleting Orai1 in cultured microglia. (*ii)* For the *in vivo* studies, we intraperitoneally injected young adult (∼5 weeks of age) mice with tamoxifen or corn oil (as controls) and employed the mice for experiments 14 days (or for the experiment in Fig. S5 34 days) later. RT-qPCR analysis of freshly isolated microglia isolated 21 days after tamoxifen administration from adult mice **(Fig. S1C)** showed marked loss of Orai1 mRNA with no change in the expression of Orai2 and Orai3 isoforms or in expression of STIM1/2, indicating that the technique provides a highly effective approach for selectively deleting Orai1 in brain microglia **(Fig. S1D)**. These microglial Orai1-specific KO cells and tamoxifen-injected mice will henceforth be referred to as mgOrai1 KO cells/mice. Orai1*^fl/fl^* mice and Orai1*^fl/fl^ ^Cx3CR1-Cre/ERT2^* mice exposed to the vehicle (corn oil) were used as controls.

Staining with the microglial marker IBA1 revealed that primary cells isolated from the spinal cord using our isolation procedures were >90% positive for IBA1 **(Fig. 1A)**, confirming the microglial identify of the cultured cells. To study the role Orai1 for mediating SOCE in microglia, we loaded cells with the ratiometric dye, Fura-2/AM and monitored cellular [Ca^2+^]_i_ elevations using a widely employed Ca^2+^-addback protocol for assessing SOCE (*42*, *46*). In this protocol, we depleted endoplasmic reticulum Ca^2+^ stores with the SERCA inhibitor, thapsigargin, administered in a Ca^2+^-free medium, and measured SOCE by examining the rate and amplitude of Ca^2+^ influx seen upon restoring extracellular Ca^2+^ (2 mM) to the bath medium **(Fig. 1B)**. WT (Orai1*^fl/fl^*) or Orai1*^fl/fl^ ^Cx3CR1-Cre/ERT2^*microglia not exposed to tamoxifen showed robust SOCE with pharmacological properties consistent with those of Orai1, including blockade by low doses of La^3+^ and BTP2 **(Figs. 1B&C)** and by the recently developed Orai1 inhibitor, CM4620, which is currently in clinical trials for treating Orai1-mediated inflammatory syndromes (*47*, *48*) **(Fig. S2A)**. SOCE was similar in magnitude in the Orai1*^fl/fl^ ^Cx3CR1-Cre/ERT2^* microglia not treated with tamoxifen compared to that seen in wild-type (Orai1*^fl/fl^*mice) **(Fig. 1B,C)**, indicating that expression of Cre recombinase alone in the absence of tamoxifen has no effects on SOCE. By contrast, Orai1*^fl/fl^ ^Cx3CR1-Cre/ERT2^* microglia exposed to tamoxifen (mgOrai1 KOs) showed marked loss of SOCE **(Fig. 1C)**, with the amplitude and the rate of Ca^2+^ entry following Ca^2+^ re-addition decreasing >90% **(Fig. 1C)**. Moreover, analysis of SOCE in cells cultured separately from individual male and female pups revealed that microglia from both sexes exhibited comparable levels of SOCE **(Fig. S2B,C)**. Importantly, deletion of Orai1 resulted in loss of SOCE in both male and female microglia **(Fig. S2B, C)** indicating that *in vitro*, there is no sex difference in the contribution of Orai1 for mediating SOCE in microglia. Microglia lacking Orai1 also displayed reduced release of Ca^2+^ from ER Ca^2+^ stores following store depletion by thapsigargin **(Fig. 1D)**, indicating that loss of SOCE affects store refilling as expected for a Ca^2+^ pathway responsible for recharging and replenishing stores. Taken together, these results indicate that Orai1 is essential for mediating SOCE in spinal cord microglia, a role congruent with the well-established role of Orai1 in many non-excitable cells (*37*, *49*).

**Figure 1.**
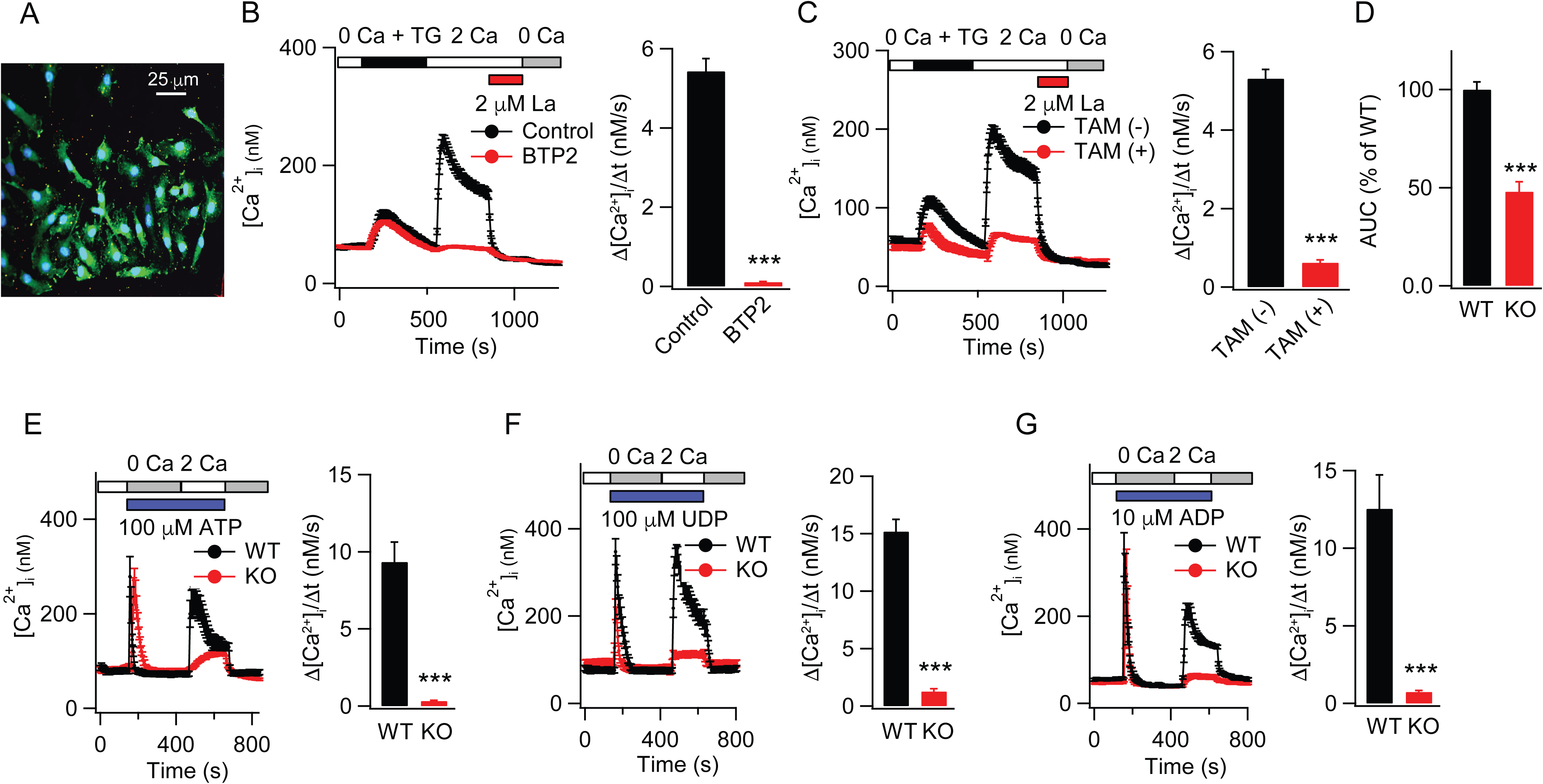
Deletion of Orai1 abrogates SOCE in spinal cord microglia. **(A)** IBA1 staining for microglia. Primary cells cultured from the spinal cord were immunolabelled with the microglial/macrophage marker, IBA1 to assess the microglial identity of the cultured cells. Blue: DAPI, Green: IBA1. **(B)** Microglia exhibit SOCE with pharmacological properties consistent with CRAC channels. Depletion of ER Ca^2+^ stores with thapsigargin (TG) in Ca^2+^-free solution evokes store release, followed by SOCE in response to addback of extracellular Ca^2+^ (2 mM). SOCE is blocked by the CRAC channel inhibitors, La^3+^ and BTP2. The bar graph on the right summarizes the rate of Ca^2+^ influx rate by measuring the initial slope of Ca^2+^ entry over 18 s after re-addition of 2 mM Ca^2+^ solution from 3-5 experiments (shown as mean±sem). n=43 cells (control), n=56 cells (BTP2)**. (C)** SOCE is abrogated by deletion of Orai1. Microglia obtained from Orai1*^fl/fl^ ^Cx3CR1-Cre/ERT2^* mice were cultured either in the absence or presence of tamoxifen. The bar graphs summarize changes in the rate of SOCE following addback of extracellular Ca^2+^ absence or presence of tamoxifen. n=56 cells (no tamoxifen), n=45 cells (tamoxifen). **(D)** Intracellular store release is diminished in Orai1 KO microglia. Store release was assessed by quantifying the area under the curve during the application of thapsigargin in the Ca^2+^-free solution until the start time of extracellular Ca^2+^ re-addition over 378 s. The mean WT levels were normalized to 100%. n=56 cells (no tamoxifen), n=45 cells (tamoxifen). **(E)** Stimulation of microglia with ATP in a Ca^2+^ - free solution evokes store release, followed by a transient SOCE when extracellular Ca^2+^ is added back. SOCE is strongly reduced in the mgOrai1 cells. n=35 cells (WT + ATP), n=50 cells (Orai1 KO + ATP). **(F,G)** Likewise, the selective P2Y receptor agonists, UDP, which stimulates P2Y_6_ receptors, and ADP, which stimulates P2Y_12_ receptors evoke SOCE which is abrogated in Orai1*^fl/fl^ ^Cx3CR1-Cre/ERT2^* microglia. Ligands were administered in Ca^2+^-free media to evoke ER Ca^2+^ release and extracellular Ca^2+^ was readded 200 s later to trigger SOCE. UDP: n=35 (WT), n=39 cells (Orai1 KO). ADP: n=48 (WT) and n=58 (Orai1 KO). In each case, the right graphs summarize the rate of Ca^2+^ influx following re-addition of extracellular Ca^2+^from three to five independent experiments. ***p<0.001 by Mann-Whitney rank sum test.

### Deletion of Orai1 impairs ATP-mediated Ca^2+^ signals in microglia

Previous studies have shown that the nucleotide neurotransmitter, ATP, which also functions as a danger-associated molecular pattern (DAMP), is a potent activator of microglia, inducing a variety of microglial responses including altered expression of inflammation-related genes, process retraction, migration, and increased phagocytic ability (*3*, *4*). Importantly, ATP released from sensory afferents and other dorsal spinal neurons following nerve injury is thought to be a key player in the activation of microglia and induction of neuropathic pain (*7*, *8*). We have recently shown that extracellular ATP is a potent activator of Orai1-mediated SOCE in astrocytes and airway epithelial cells (*42*, *50*) raising the possibility that Orai1 channels may similarly mediate ATP-induced Ca^2+^ responses in microglia. To directly address this question, we examined Ca^2+^ signals evoked by agonists of ATP receptors. We found that administration ATP in Ca^2+^ -free medium evoked robust, transient rises in [Ca^2+^]_i_ in WT microglia reflecting release of Ca^2+^ from intracellular stores **(Fig. 1E)**. Re-addition of extracellular Ca^2+^ evoked a robust secondary rise in [Ca^2+^]. The kinetic signature of this ATP-induced Ca^2+^ response is consistent with Gq/phospholipase C-inositol 1,4,5-triphosphate signaling and store-operated calcium entry (SOCE) that we and others have previously described in AECs and other cell types for P2Y receptors (*42*, *50*). In agreement with this interpretation, UDP, a uridine nucleotide agonist of the P2Y6 receptor, and 2-methylthio-ADP (2-MeSADP or ADP), a ligand for the P2Y12 receptor which are both strongly implicated microglia-mediated inflammation and neuropathic pain (*9*-*11*, *39*) also evoked Ca^2+^ elevations with amplitude and kinetics comparable to those evoked by ATP **(Fig. 1F,G)**. Thus, these results indicate that ATP stimulates Ca^2+^ signaling in microglia at least in part through the activation of metabotropic purinergic receptors.

To examine the role of Orai1 in these Ca^2+^ responses, we examined mgOrai1 KO cells in the same protocol. Deletion of Orai1 caused near complete loss of SOCE evoked by ATP **(Fig. 1E)**. Likewise, deletion of Orai1 impaired SOCE evoked by UDP **(Fig. 1F)** and by 2-MeSADP (ADP) **(Fig. 1G)**. Together, these results, indicate that CRAC channels composed of Orai1 are essential for the SOCE that is activated by stimulation of P2Y6 and P2Y12 receptors in spinal microglia.

Another receptor that is implicated in microglial activation and chronic pain is TLR4. Intrathecal injection of the endotoxin lipopolysaccharide (LPS), a potent activator of TLR4 causes symptoms of neuropathic pain (*51*, *52*) and increasing evidence suggests that genetic or pharmacological blockade of TLR4 in pain models protects against neuroinflammation and hyperalgesia is typically seen following nerve injury (*51*-*54*). Although Ca^2+^ mobilization downstream of TLR4 activation is not widely reported, LPS is known to activate SOCE and CRAC currents in mouse cortical microglia (*55*). We found that administration of LPS elicited an initial Ca^2+^ rise followed by slowly occurring Ca^2+^ oscillations in WT microglia that lasted for > 1 hour **(Fig. S2D)**. However, in mgOrai1 KO microglia, these low-frequency LPS-evoked Ca^2+^ fluctuations were completely abrogated, indicating that SOCE through Orai1 contributes to endotoxin-mediated Ca^2+^ signals in microglia. Taken together, these results indicate that Orai1 channels are a major mechanism for mobilization of Ca^2+^ influx in microglia by a variety of receptors implicated in neuropathic pain.

### Orai1 promotes production of pro-inflammatory cytokines from spinal microglia

Microglia are known for their prolific production and release of proinflammatory cytokines which are thought to drive neuroinflammation in disease states. Given the strong links between Orai1-mediated Ca^2+^ influx and Ca^2+^-dependent gene expression of cytokines in many cells (*56*), we next sought to determine if Orai1 has a role in this function of microglia. We addressed this question by activating microglia with LPS, a widely used stimulus for inducing microgliosis and microglial-mediated cytokine production, or by exposing cells to thapsigargin to directly deplete Ca^2+^ stores. In line with previous evidence (*57*, *58*), we found that stimulating microglia with LPS caused strong induction of the proinflammatory mediators IL-6, TNF⍺, MCP1, and PGE_2_ **(Fig. 2B-D)**. In Orai1 KO microglia, however, LPS-mediated induction of these proinflammatory mediators was markedly reduced **(Fig. 2B-D)**. By contrast, IL-10, an anti-inflammatory cytokine, was unaffected by LPS and unchanged by deletion of Orai1 suggesting that Orai1 has a stronger role in stimulating inflammatory cytokines **(Fig. S3)**. Similarly, deletion of Orai1 also strongly impaired IL-6 and TNF⍺ production from primary spinal microglia stimulated with TG+PdBu to directly evoke SOCE **(Fig. 2E,F).** Moreover, detailed examination of these cytokines in microglia cultured separately from male and female mice, showed that both the degree of induction of IL-6, TNF⍺, MCP1, and PGE_2_ as well as the effects of knocking out Orai1 were comparable between isolated from male and female mice (**Fig. 2**). Together, these *in vitro* results indicate that Orai1 plays a key role in the synthesis and release of proinflammatory cytokines from spinal microglia.

**Figure 2.**
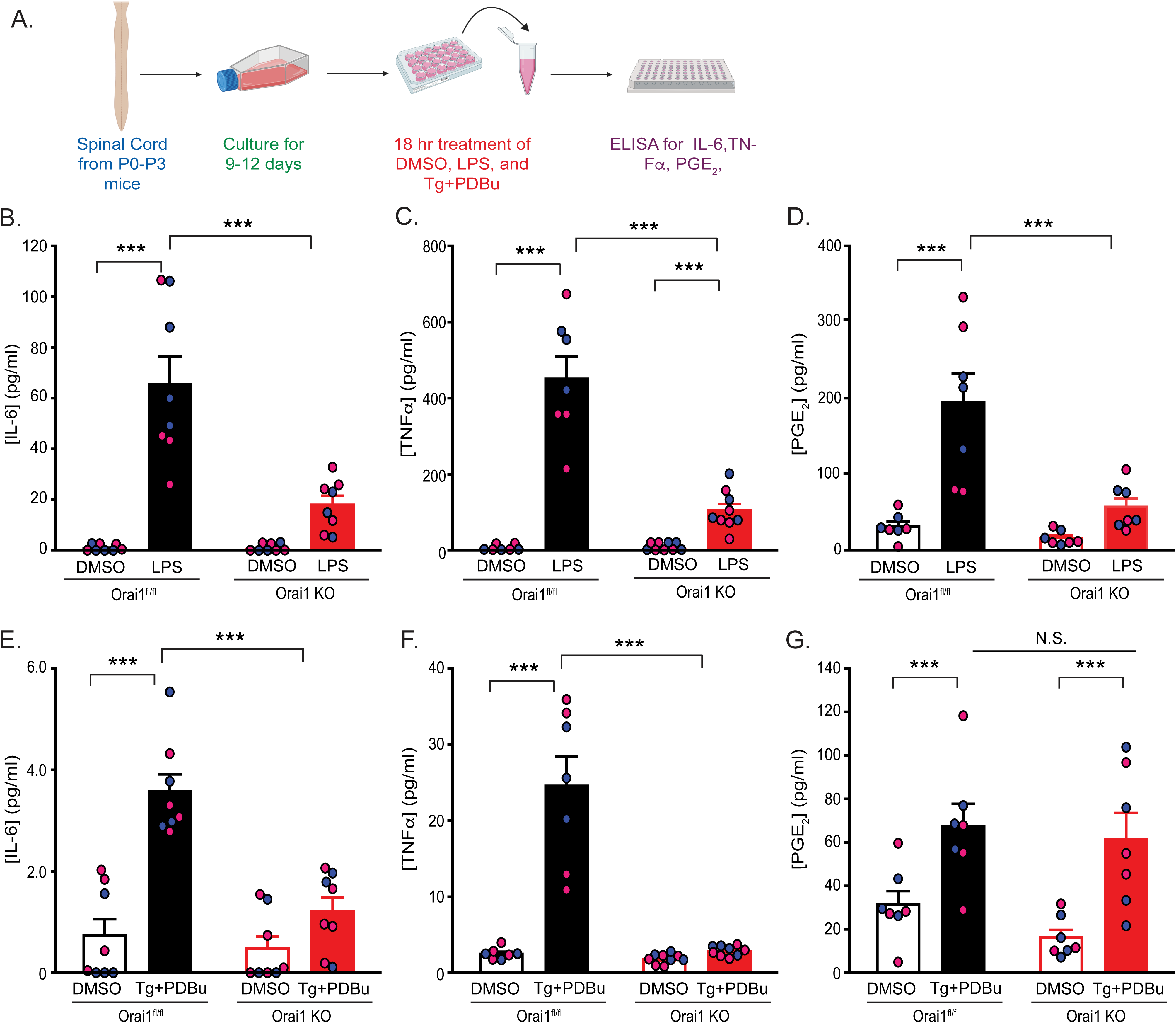
Deletion of Orai1 impairs synthesis of proinflammatory mediators from spinal microglia. **(A-D)** WT (Orai1*^fl/fl^*) or mgOrai1 KO primary microglia cultured for 7-10 days and then stimulated with LPS (1 ng/ml) for 18 hours. Levels of IL-6, TNF⍺, and the lipid mediator, PGE_2_ were measured in the cell culture supernatant by ELISA. Cells obtained from male and female pups were cultured separately. Due to the low yield of cells, cells from two pups of the same sex were combined prior to the experiment. Therefore, each point represents the measurement from one flask (two animals). n=3-7 measurements from 6-14 mice. ***: p<0.001 by two-way ANOVA followed by Tukey test for comparison between multiple groups. **(E-H)** Cytokine measurements in microglia stimulated with thapsigargin and PdBu. Cytokine levels were measured in the supernatant 18 hours following cell stimulation. N=3-7 measurements from 6-14 mice. ***: p<0.001 by two-way ANOVA followed by Tukey test for comparison between multiple groups.

### Ablation of Orai1 in microglia protects male mice against neuropathic pain

Inflammatory cytokines such as TNF⍺ and IL-6 are strongly implicated in the etiology of neuropathic pain (*20*, *21*, *59*). Thus, the finding that deletion of Orai1 reduces the production of proinflammatory cytokines from spinal microglial led us to consider the consequences for neuropathic pain. We examined this question by assessing allodynia in WT and mgOrai1 KO mice following peripheral nerve injury **(Fig. 3A)**. We used the spared nerve injury (SNI) model to induce peripheral nerve injury in young adult mice (∼6 weeks), a model in which the sciatic nerve is exposed at the trifurcation of the sural, tibial, and common peroneal nerves, and the tibial and common peroneal nerves are ligated and severed, leaving the sural nerve intact (*60*, *61*). SNI is a robust model of neuropathic pain where every animal with SNI exhibits mechanical and cold allodynia for the remainder of animal life (*61*). Paw withdrawal thresholds to Von Frey filament (VF) stimulation applied every 2-3 days were used to assess mechanical sensitivity of the hind paws before and following the SNI surgery for 14 days as previously described (*60*). All pain-related behaviors were assessed using double-blind procedures.

**Figure 3.**
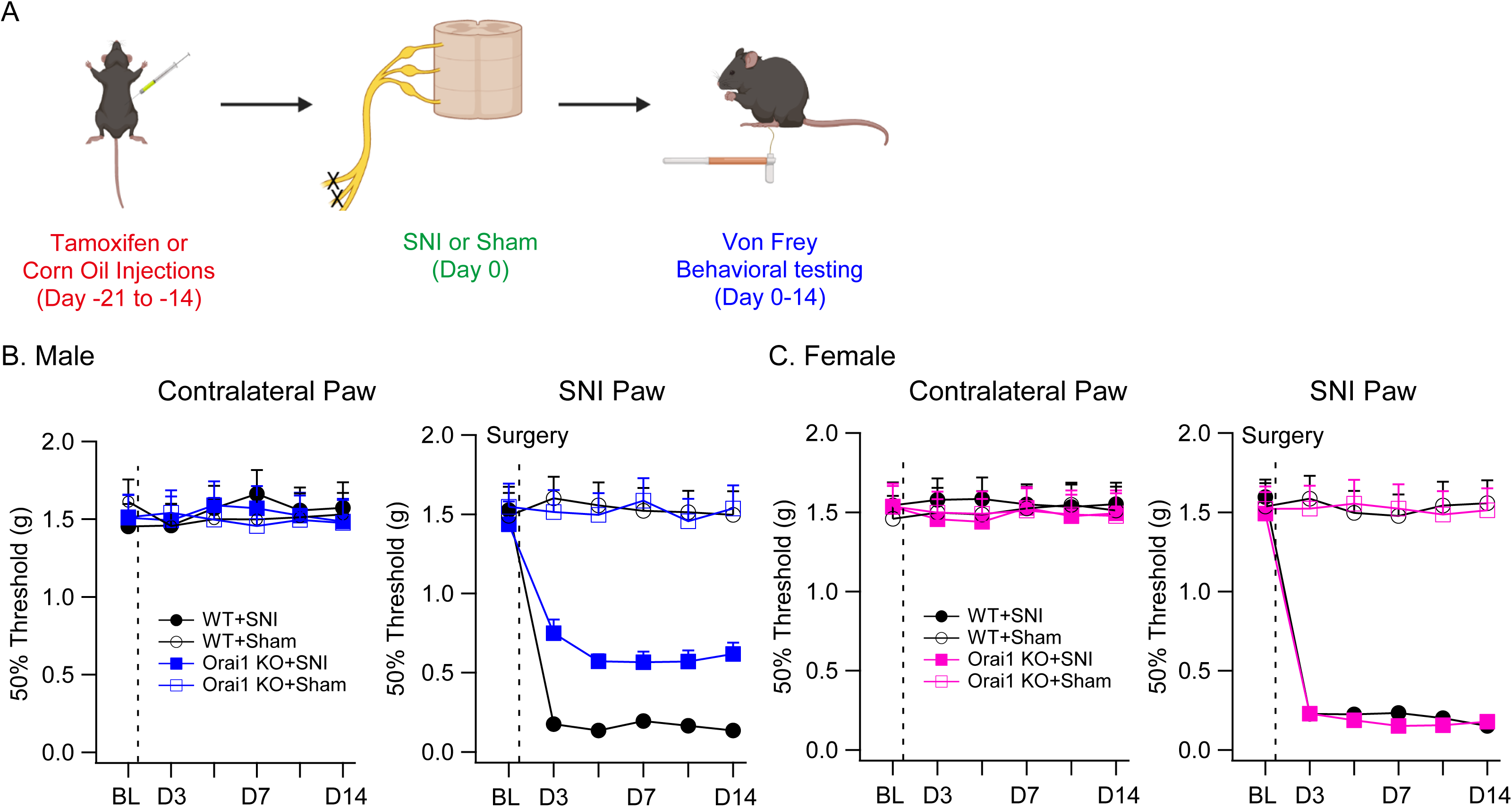
Male microglial Orai1 KO mice are partially protected in the SNI model of neuropathic pain. **(A)** A schematic of the experiment. WT (Orai1 *^fl/fl^*) or mgOrai1 KO (Orai1*^fl/fl^ ^Cx3CR1-Cre/ERT2^* with tamoxifen) mice (8 weeks old) were subjected to SNI or sham surgery on the left sciatic nerve, and mechanical sensitivity measured by von Frey thresholds were monitored at baseline (prior to SNI) and for 14 days following SNI surgery. **(B,C)** von Frey thresholds in male

*(B)* or female *(C)* mice before and after SNI surgery in the indicated genetic groups and conditions. The dotted line indicates the day of SNI surgery. N values are as follows: Male WT mice: n=10; Male Orai1 KO mice: n=10; Female WT mice: n=11; Female Orai1 KO mice: n=9. *: p<0.05 by unpaired *t*-test for each time point comparing WT and mgOrai1 KO mice subjected to SNI.

SNI caused paw withdrawal thresholds to markedly decrease on days 3-14 following the nerve injury **(Fig. 3)**. Sham-operated mice did not show the hypersensitivity in line with previous studies indicating the nerve injury causes robust allodynia in this model of neuropathic pain (*61*). mgOrai1 KO mice showed comparable paw withdrawal thresholds as WT mice prior to SNI, suggesting that microglial Orai1 channels do not directly regulate mechanical sensation. Following SNI, however, male mgOrai1 KO mice showed significant protection against allodynia compared to sham-operated or WT (Orai1*^fl/fl^*) mice **(Fig. 3B)**. Mitigation of allodynia in the mgOrai1 KO male mice was robust, lasting >14 days following SNI surgery. Orai1*^fl/fl^ ^CX3CR1-Cre/ERT2^*mice exposed only to the vehicle (corn oil) did not show reduction in allodynia following SNI, in contrast to littermate male Orai1*^fl/fl^ ^CX3CR1-Cre/ERT2^* mice injected with tamoxifen **(Fig. S4)**, thus ruling out Cre-dependent effects in the mitigation of allodynia.

One concern in the above experiments is that because CX3CR1 is also expressed in peripheral inflammatory macrophages and monocytes, a potential role for infiltrating macrophages with loss of Orai1 in the mitigation of the pain phenotype cannot be formally ruled out. This possibility is moderated by the fact that monocytes and circulating macrophages are relatively short-lived and turn over rapidly in a few days (*62*-*64*)) whereas resident microglia turn-over extremely slowly and are further largely self-renewing (*62*, *65*, *66*). Moreover, reporter labelling, studies have shown that CX3CR1-Cre mice when used 30 days or later after tamoxifen permit Cre-dependent recombination almost exclusively in microglia (*62*). Additionally, Renthal and colleagues have found undetectable levels of Orai1 transcript in DRG macrophages (*67*). Because we observed stable and robust pain mitigation in Orai1*^fl/fl^ ^CX3CR1-Cre^*male mice even 28 days after the last tamoxifen pulse **(Fig. 3B)**, it is unlikely that the observed phenotype is due to peripheral monocytes lacking Orai1. However, to address this concern further, we increased the duration of the waiting period before nerve injury for recombination after the final tamoxifen injection from 14 to 34 days by which time peripheral CX3CR1^+^ cell populations should be replaced by non-recombined cells from wildtype progenitors. We then assessed von Frey paw withdrawal thresholds for 18 days after SNI (i.e., 52 days after the final tamoxifen injection). These experiments also showed highly significant pain mitigation in the Orai1*^fl/fl^ ^CX3CR1-Cre^* male mice on the injured side **(Fig. S5)**. In fact, the degree of pain mitigation in mgOrai1 KO mice was comparable to that seen in the cohort of mice subjected to the shorter waiting duration **(Fig 3 vs Fig. S5)**. Thus, the most straightforward interpretation of these behavioral tests is that the loss of pain hypersensensitivity in male Orai1*^fl/fl^ ^CX3CR1-Cre^* mice is due to selective deletion of Orai1 in microglia.

In contrast to male mice, we were surprised to see that female mice behaved no differently than WT control mice **(Fig. 3C)**. In this group, paw withdrawal thresholds after SNI in female mgOrai1 KO mice were not different from those seen in Orai1*^fl/fl^* mice **(Fig. 3C)**. Thus, protection against SNI-induced neuropathic pain in mgOrai1 KO mice is highly sex-specific. The finding that deletion of Orai1 in microglia produces a gender-specific phenotype similar to that previously described with microglial blockers or microglial poisoning (*23*) suggests that the gender-effect is traceable to a specific signaling pathway in microglia rather than differences in downstream neural circuits in the CNS.

### Deletion of Orai1 reduces spinal cytokine levels *in vivo* in response to nerve injury

It is well-established that microglia produce inflammatory cytokines that drive inflammation mediating neuropathic pain. Given that our *in vitro* experiments indicated that deletion of Orai1 blocked the induction of key cytokines in cultured microglia, we next considered whether the partial mitigation of mechanical allodynia in male mgOrai1 KO mice is due to reduction in cytokine levels *in vivo* in the spinal cord following SNI. To address this question, we assessed cytokine levels in the spinal cord lumbar tissue in WT and mgOrai1 KO mice following peripheral nerve injury. The lumbar spinal cord (5 mm in length) at L4-5 encompassing the entry of the sciatic nerve was dissected out following SNI or sham surgery from Orai1*^fl/fl^* and Orai1*^fl/fl^ ^CXCR3-Cre/ERT2^* mice treated with tamoxifen (*68*, *69*), and cytokine levels in the homogenized tissue were measured via ELISA as described in previous reports (*68*, *69*) **(Fig. 4)**. These measurements revealed that SNI evokes marked increases in TNFα, IL-6, IL1-β, and BDNF in the spinal cords of WT mice compared to sham-operated controls **(Fig. 4B-E)**. Both male and female WT mice showed comparable increases in spinal cytokine levels relative to sham-operated controls following nerve injury **(Fig. 4)**. Strikingly, however, the concentrations of these inflammatory cytokines were significantly reduced in male mgOrai1 mice, with IL-6 in particular declining essentially to levels found in sham-operated controls **(Fig. 4)** (154±10 pg/ml in SNI WT mice versus 72±7 pg/ml in SNI mgOrai1 KO mice; p<0.001). Levels of TNFα, IL1-β, and BDNF also decreased in mgOrai1 KO mice relative to WT mice following SNI **(Fig. 4C,D,E)**. However, female mgOrai1 KO mice subjected to SNI did not show reductions of these mediators **(Fig. 4B-E)**. Rather, the degree of induction of IL-6, TNFα, BDNF, and IL1-β in female Orai1 KO mice was comparable to WT mice with SNI. These results indicate that deletion of microglial Orai1 channels prevents the induction of key proinflammatory cytokines in the spinal cord that are implicated in neuropathic pain following nerve injury, but this is seen only in male mice.

**Figure 4.**
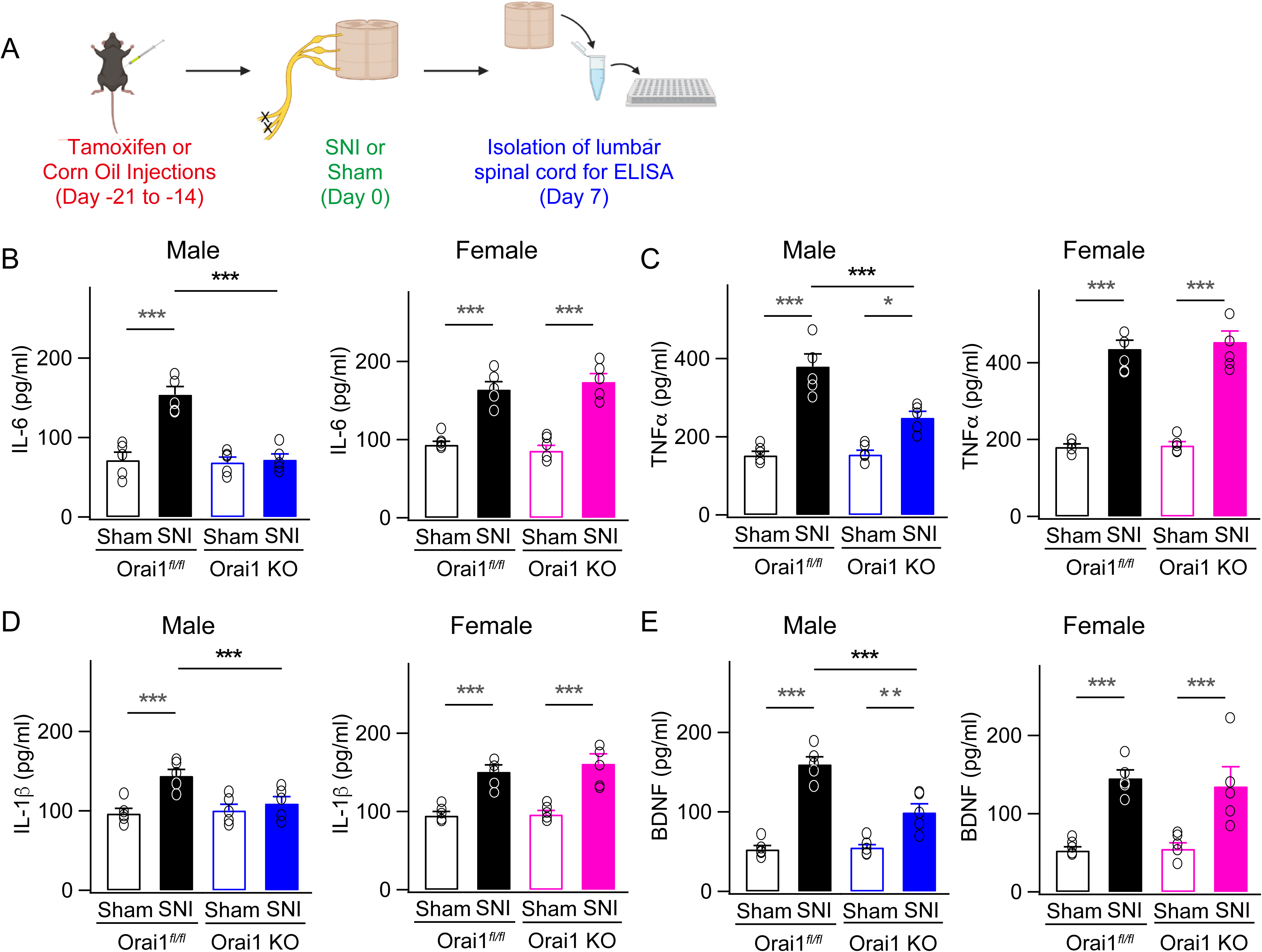
Deletion of Orai1 in microglia blocks increases in proalgesic cytokines in the spinal cord following SNI. **(A)** A schematic of the protocol used for the experiment. Wildtype (Orai1*^fl/fl^*) or mgOrai1 KO (Orai1*^fl/fl^ ^CX3CR1-CRE/ERT2^* with tamoxifen) mice were subjected to either sham or SNI surgery. 7 days following SNI, animals were euthanized, 5 mm of the lumbar spinal cord was harvested from each mouse, and the tissue was homogenized and spun down. Levels of IL-1β, TNF⍺, IL-6, and the growth factor BDNF were assessed in the supernatant using ELISA. **(B-I)** SNI causes increases in the levels of inflammatory cytokines and the growth factor BDNF compared to sham-operated controls. These increases are reduced in male but not female mgOrai1 KO mice. (N=5 mice per condition). ***: p<0.001; *: p<0.05 by one-way ANOVA followed by Tukey test for comparison between multiple groups.

### SNI-induced enhancement of glutamatergic neurotransmission in the spinal cord is partially occluded in male mgOrai1 KO mice

Previous work has shown that peripheral nerve injury and microglial stimulation increases excitatory synaptic transmission between lamina II neurons of the dorsal spinal cord (*59*, *70*-*76*). This maladaptive potentiation of excitatory synaptic transmission is thought to alter the set-point for signal transmission in nociceptive circuits following nerve injury to mediate neuropathic pain (*4*, *14*, *16*). Thus, the finding that male mgOrai1 KO mice are protected against SNI-induced allodynia **(Fig. 3B)** with reduced spinal cytokine levels **(Fig. 4)** led us to next consider whether microglial Orai1 channels regulate SNI-evoked alterations in synaptic transmission in the spinal cord. To address this question, we carried out electrophysiological analysis of spontaneous excitatory postsynaptic currents (sEPSCs) in the substantia gelatinosa (SG) neurons of the dorsal spinal cord (L4-5) following SNI **(Fig. 5A)**. The SG region of the dorsal horn was visually identified in spinal cord slices with a water immersion objective, and sEPSCs were recorded from SG neurons in the whole-cell patch-clamp mode at -70 mV in the presence of GABA and NMDAR blockers as described previously (*43*, *77*).

**Figure 5.**
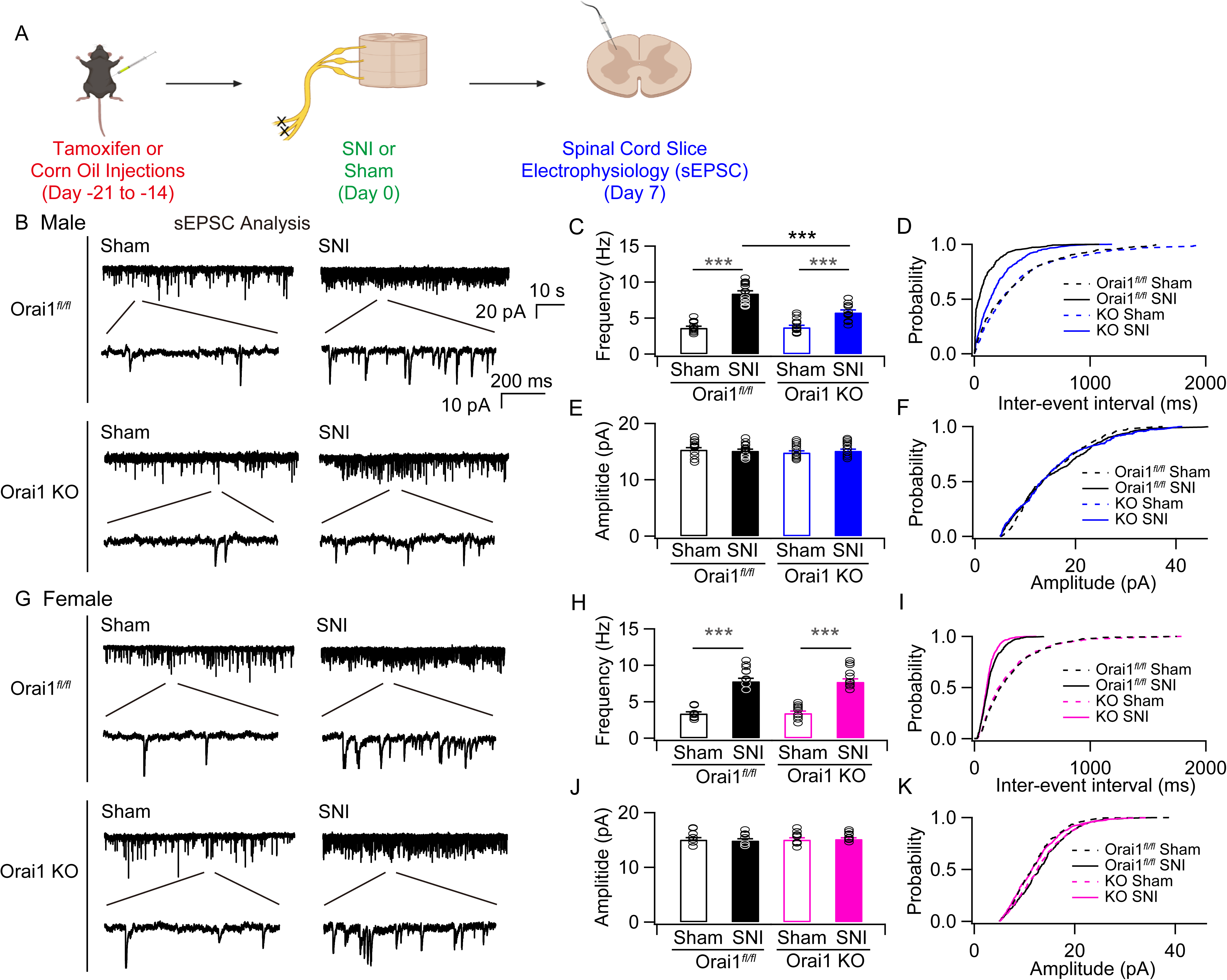
SNI-induced increase in the frequency of sEPSCs in the dorsal horn spinal neurons is mitigated in male mgOrai1 KO mice. **(A)** A schematic of the experiment. WT (Orai1 *^fl/fl^*) or mgOrai1 KO (Orai1*^fl/fl^ ^Cx3CR1-Cre/ERT2^* with tamoxifen) mice (8 weeks old) were subjected to SNI or sham surgery. 7-8 days following SNI, animals were euthanized and the spinal cords isolated for slice electrophysiology. **(B)** sEPSCs recorded in the whole-cell patch-clamp configuration (-70 mV) from a lamina II neurons in the dorsal horn of the L4 spinal cord in slices from male mice. Ionotropic GABA receptors and NMDARs were blocked with picrotoxin and D-APV. The traces show examples of sEPSCs at low and high time resolutions. **(C)** Bar graph summarizing the frequency of sEPSCs in the indicated groups. Mice subjected to SNI show increase in sEPSCs which is partially occluded in mgOrai1 mice. ***: p<0.001. One-way ANOVA followed by Tukey test for comparison between multiple groups. **(D)** Cumulative probability distributions of the inter-event intervals of the sEPSCs in WT and mgOrai1 KO mice. The inter-event interval is shifted towards lower intervals consistent with increases in the frequency of sEPSCs in mice subjected to SNI. **(E)** Summary of the sEPSC amplitudes in the indicated groups. There is no change in the amplitude of the events. **(F)** Cumulative distributions of the sEPSC amplitudes. **(G)** sEPSC traces in slices from female mice. **(H,I)** Summary of the changes in mean frequency and cumulative histogram distributions of sEPSC frequency in female mice. In female mgOrai1 KO mice, the frequency of sEPSC is unaffected relative to SNI-administered WT mice. **(J-K)** Mean and cumulative probability distributions of the sEPSC amplitudes. N values are as follows: Male WT mice: n=12 cells (sham), n=13 (SNI). Male Orai1 KO mice: n=13 cells (sham), n=13 cells (SNI). Female WT mice: n=10 cells (sham), n=10 cells (SNI). Female Orai1 KO mice: n=10 cells (sham), n=10 cells (SNI). ***: p<0.001 by ANOVA followed by Tukey test for comparison of multiple groups.

Neuronal recordings from sham-operated WT and mgOrai1 KO slices revealed no differences in the frequency or amplitude of basal sEPSC events between the two genetic groups **(Figs. 5C,E and 5H,J)**. However, WT animals that underwent SNI exhibited striking increases in the frequency of sEPCSs, with the frequency increasing approximately two-fold in SNI-administered WT mice (from 3.6±0.2 Hz (n=12) in sham-operated animals to 8.4±0.38 Hz (n=13) (p<0.001) in SNI animals) **(Fig. 5C)**. Cumulative histogram plots indicated that SNI significantly decreased the inter-event sEPSC interval **(Fig. 5D)**, consistent with the increase in the frequency of sEPSCs. No change in the amplitude of the events, however, was detected **(Fig. 5E,F)**. These changes are consistent with previous literature indicating that peripheral nerve injury increases spontaneous excitatory synaptic transmission in the lamina II neurons of the dorsal spinal cord (*71*, *74*-*76*, *78*).

In contrast to the >2-fold increase in sEPSCs frequency seen in WT mice following SNI, male mgOrai1 KO subjected to SNI showed significantly smaller potentiation of the sEPSC frequency (8.4±0.38 Hz (n=13) in sham-operated KOs versus 5.7±0.3 (n=11) in SNI animals, p<0.001) **(Fig. 5C)**. Correspondingly, mgOrai1 KO mice also showed smaller changes in inter-event sEPSC intervals **(Fig. 5D)**. These results indicate that deletion of Orai1 in microglia partially prevents SNI-induced enhancement of excitatory synaptic transmission in lamina II dorsal horn neurons in male mice.

In female mice, sEPSCs in slices from WT (Orai1^fl/fl^) mice subjected to SNI also showed increases in the frequency of sEPSCs (from 3.4±0.2 Hz (n=8) in sham-operated animals to 7.8±0.4 Hz (n=9) (p<0.001) in SNI animals) (Figure 3H,I). However, in contrast to male mice, female mgOrai1 KO mice subjected to SNI did not show attenuation of the sEPSC frequency (3.5±0.2 Hz (n=9) in sham-operated and 7.7±0.4 Hz (n=10) (p<0.001) in SNI animals), and the extent of EPSC potentiation remained similar to that seen in the female WT (Orai1*^fl/fl^*) counterparts **(Fig. 5H,I)**. Thus, in female mice, deletion of microglial Orai1 does not dampen the maladaptive synaptic potentiation in response to nerve injury.

Because changes in inhibitory synaptic transmission are also thought to contribute to chronic pain (*4*, *16*), we next examined if Orai1 modulates inhibitory synaptic transmission following nerve injury. In contrast to the changes in excitatory synaptic transmission, we did not observe alterations in the frequency of spontaneous inhibitory synaptic currents (sIPSCs) **(Fig. S6)**, nor were there any changes in the amplitude of the IPSCs following SNI **(Fig. S6)**. Likewise, SNI did not induce alterations in the amplitude or frequency of the miniature IPSCs **(Fig. S7)**. Further, no differences in either frequency or amplitude were seen between the WT and mgOrai1 KO groups (e.g., **Fig. S7)**. Thus, under our experimental conditions, SNI did not elicit changes in the frequency or amplitude of inhibitory synaptic transmission in lamina II neurons in the mouse spinal cord.

To further assess the mechanisms of synaptic potentiation by SNI and the contributions of microglial Orai1 channels to this process, we carried out measurements of miniature EPSCs (mEPSCs) in the absence of action potentials to address potential changes in the presynaptic release probability **(Fig. S8)**. As shown previously (*79*), we found that in the presence of 1 µM TTX to block action potentials, SNI-treated mice showed significant increases in the frequency of mEPSCs **(Fig. S8C)**. Analysis of the miniature events revealed significant shortening of the inter-event intervals **(Fig. S8)**, indicating that nerve injury enhances the rate of glutamate release from excitatory lamina II neurons. Male Orai1*^CX3CR1-Cre/ERT2^* mice with deletion of Orai1 in microglia showed smaller enhancement in the mEPSC frequency and shortening of the inter-event intervals compared to WT controls **(Fig. S8C,D)**. As seen for spontaneous EPSCs, this phenotype in the mEPSC frequency was only seen in male mice. In female mice, there was no effect of ablating Orai1 on the mEPSC frequency or inter-event intervals **(Fig. S8H,I)**.

Taken together, these results indicate that Orai1 channels in microglia at least partially drive the SNI-mediated increase in presynaptic glutamate release probability in male mice, which contributes to the overall increase in excitatory synaptic transmission in the spinal cord following nerve injury. In female mice, by contrast, loss of microglia Orai1 channels has no effect in mitigating the maladaptive potentiation of excitatory synaptic potentiation seen following nerve injury. Thus, the contribution of microglial Orai1 channels to the potentiation of excitatory synaptic transmission in response to nerve injury evidently differ between male and female mice.

#### Orai1 regulates SNI-induced microglial proliferation in the lumbar spinal cord of male mice

The finding that Orai1 deletion strongly inhibits the induction of IL-6, TNFα, and other mediators linked to neuropathic pain led us to consider the implications *in vivo* for microgliosis in the spinal cord following nerve injury. A widely used approach for assessing microgial reactivity and proliferation is through the expression of the microglial marker, IBA1. Previous studies have shown that IBA1 expression markedly increases in the spinal cord following nerve injury (*23*, *27*) indicative of increased microgliosis and proliferation. In line with these findings, we observed that IBA1 staining significantly increased in the dorsal horn of L4-L5 spinal cord sections of WT mice following SNI **(Fig. 6A)**. There was no statistically significant sex difference in the degree of IBA1 upregulation between male and female mice **(Fig. 6B,C**, p=0.54137 comparing WT SNI male vs WT SNI female groups) consistent with previous findings that microglial proliferation is comparable following SNI in both sexes (*27*). The increase in IBA1 staining was seen in the L4-5 spinal cord region but not in spinal sections away from the sciatic nerve innervation site (T13 sections), indicating that SNI of the sciatic nerve selectively increases microglial reactivity only in those regions of the spinal cord innervated by the injured nerve **(Fig S9)**. Importantly, although male mgOrai1 KO mice also showed upregulation of IBA1 following SNI, the degree of IBA1 upregulation was significantly reduced compared to WT counterparts **(Fig. 6A,B)**. This result indicates that deletion of Orai1 impairs microgliosis *in vivo* in male mice. In contrast to male mice however, female mice showed no quantitative difference in IBA1 labelling between Orai1 KO and WT groups **(Fig. 6A,C)**. Thus, deletion of Orai1 does not affect microglial proliferation in female mice. The lack of effect on microglial reactivity parallels the lack of effects on spinal cytokine levels (Fig. 4), neuropathic pain measures (Fig. 3), and degree of synaptic potentiation (Fig. 5) in the female mice suggesting a link between the differential effects of Orai1 on microgliosis and downstream effectors phenotypes in male and female mice. Taken together, these results indicate that Orai1 deletion impairs microglial activation in male but not female mice.

**Figure 6.**
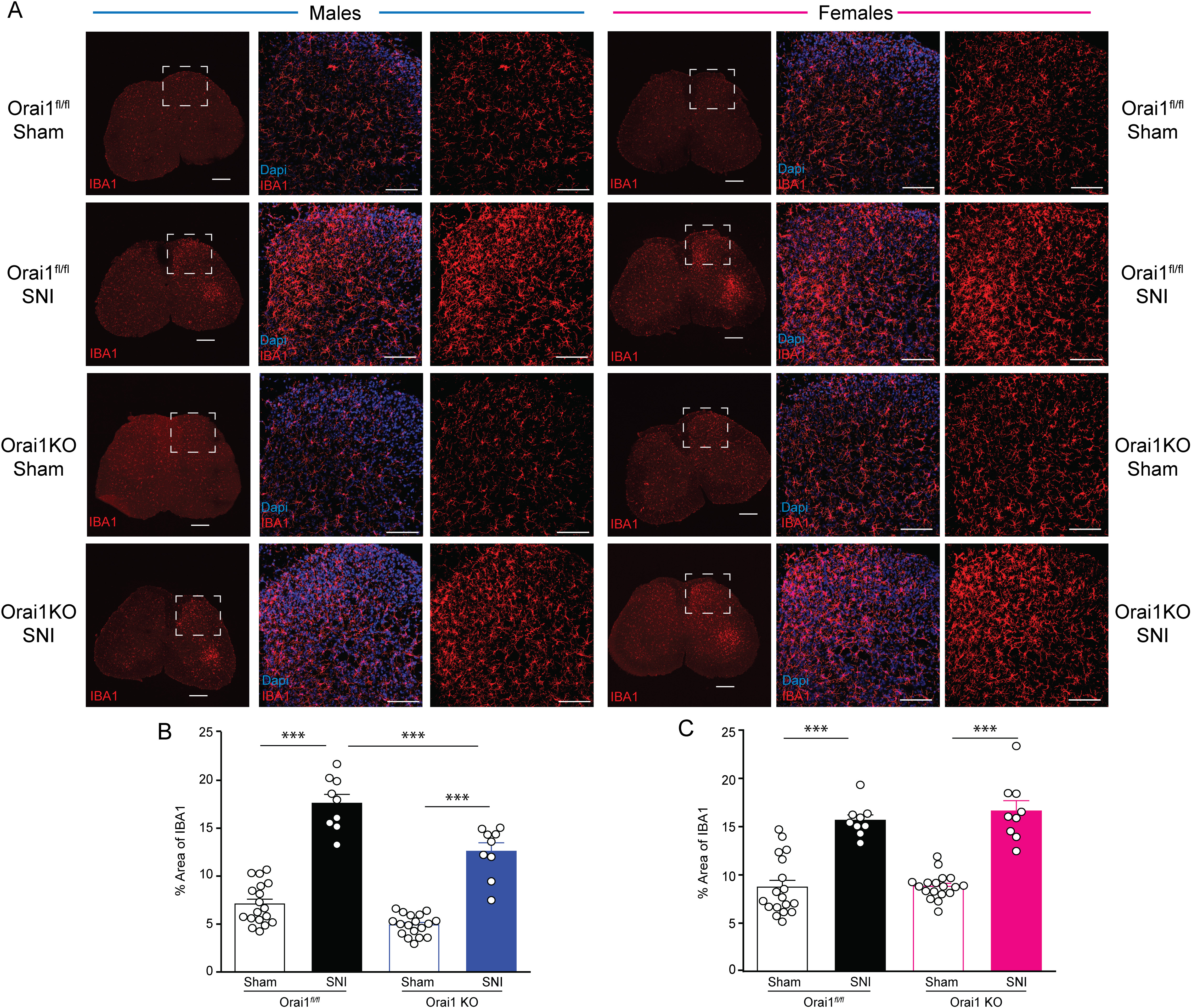
Microglial proliferation is impaired in male but not female mgOrai1 KO mice after nerve injury. **(A)** IBA1 staining of transverse sections of the lumbar spinal cord. The images in the first column (for males and females, respectively) show fixed sections of the whole spinal cord stained for IBA1. Scale bar = 250 µm. Images in the middle column show enlarged images of the dorsal horn of the lumbar spinal cord labelled with IBA1(red) and Dapi (blue). The images in the right column show the same sections with only IBA1 labelling. Scale bar = 100 µm. The dynamic range (image brightness and grey scale values) of the dorsal horn images are the same for all images displayed here. **(B,C)** Quantification of IBA1 expression in male and female mice determined by measuring the % area of the region of interest occupied by IBA1 fluorescent signal. Data are given as means +/-SEM for n = 3 images per region from 3 mice per group. (***= p<0.001 by two way ANOVA followed by Tukey between the indicated groups).

Is the upregulation in IBA1 labelling induced by SNI due to altered Orai1 expression? Despite the evidence for involvement of Orai1 in mitigating pain hypersensitivity and reducing microglial reactivity in male mice, RNA-seq analysis of the lumbar spinal cord (*80*) indicated that SNI did not evoke changes in the expression of Orai1 nor in the expression of other SOCE molecules including Orai2, STIM1, and STIM2 **(Figure S10)**. Similarly, no change in expression of the SOCE transcriptome was detected in female mice **(Figure S10)**. Thus, the most straightforward interpretation is that Orai1-mediated microgliosis in male mice following SNI is not caused by changes in Orai1 expression but instead likely mediated by the activity of existing microglial Orai1 channels.

### An Orai1 channel blocker mitigates allodynia in male but not female mice

The finding that male microglial Orai1 KO mice show attenuation of mechanical hypersensitivity following nerve injury led us to next consider whether pharmacological suppression of Orai1 can elicit a similar effect. A previous study showed that systemic delivery of the Orai1 channel antagonist, BTP2 decreases thermal and mechanical hypersensitivity following nerve injury in C57Bl/6 mice (only male mice were used in that study) (*81*). However, because BTP2 has also been shown to affect the activity of other ion channels, most notably, TRPM4 (*82*) and ryanodine receptors (*83*), the specificity of this effect is not well-resolved. We therefore sought to examine the effects of a more selective Orai1 antagonist, CM4620, which, as shown above strongly inhibits SOCE in primary cultured spinal microglia **(Fig S2A)**. CM4620 is currently in human clinical trials for mitigating inflammation in acute pancreatitis (*47*) and in patients with COVID-19 pneumonia (*48*). CM4620 shows greater selectivity for Orai1 than Orai2 or Orai3 **(Fig. S11)** and readily accumulates in the brain (see Methods). We administered CM4620 (or vehicle) by oral gavage to wildtype C57BL6 mice for 7 days beginning with the day of the SNI surgery, and assessed neuropathic pain due to nerve injury by measuring paw withdrawal thresholds to von Frey filaments as described earlier.

We found that male WT mice that received the drug were protected for mechnical hypersensitivity on days 3-7 following the SNI procedure **(Fig. 7B)**. Further, in male mice, the mitigation of allodynia only lasted for the duration of the drug administration. Cessation of the drug on day 7 caused the paw withdrawal thresholds in WT mice to return to levels seen in control mice by day 12 **(Fig. 7B**, *right graph*). This result suggests that CM4620 fundamentally does not block the induction of neuropathic pain but instead reverses the pain hypersensitivity. Female mice did not show attenuation relative to the vehicle control group **(Fig. 7C)**. To directly determine if CM4620 is also effective in relieving allodynia once it is already induced, we examined the effects of readministering the drug on day 22 after SNI (after collecting a new baseline the prior day, **Fig. 7D)**. Administering CM4620 on day 22 after the induction of mechanical allodynia also relieved hypersensitivity in male mice receiving the drug **(Fig. 7D**, *right graph*). As before, the pain relief subsided quickly (∼2 days) following cessation of the drug, similar to the rapid development of allodynia when the drug was applied immediately with nerve injury **(Fig. 7B)**. Female mice, however, did not show any attenuation of the hypersensitivity **(Fig. 7E)**.

**Figure 7.**
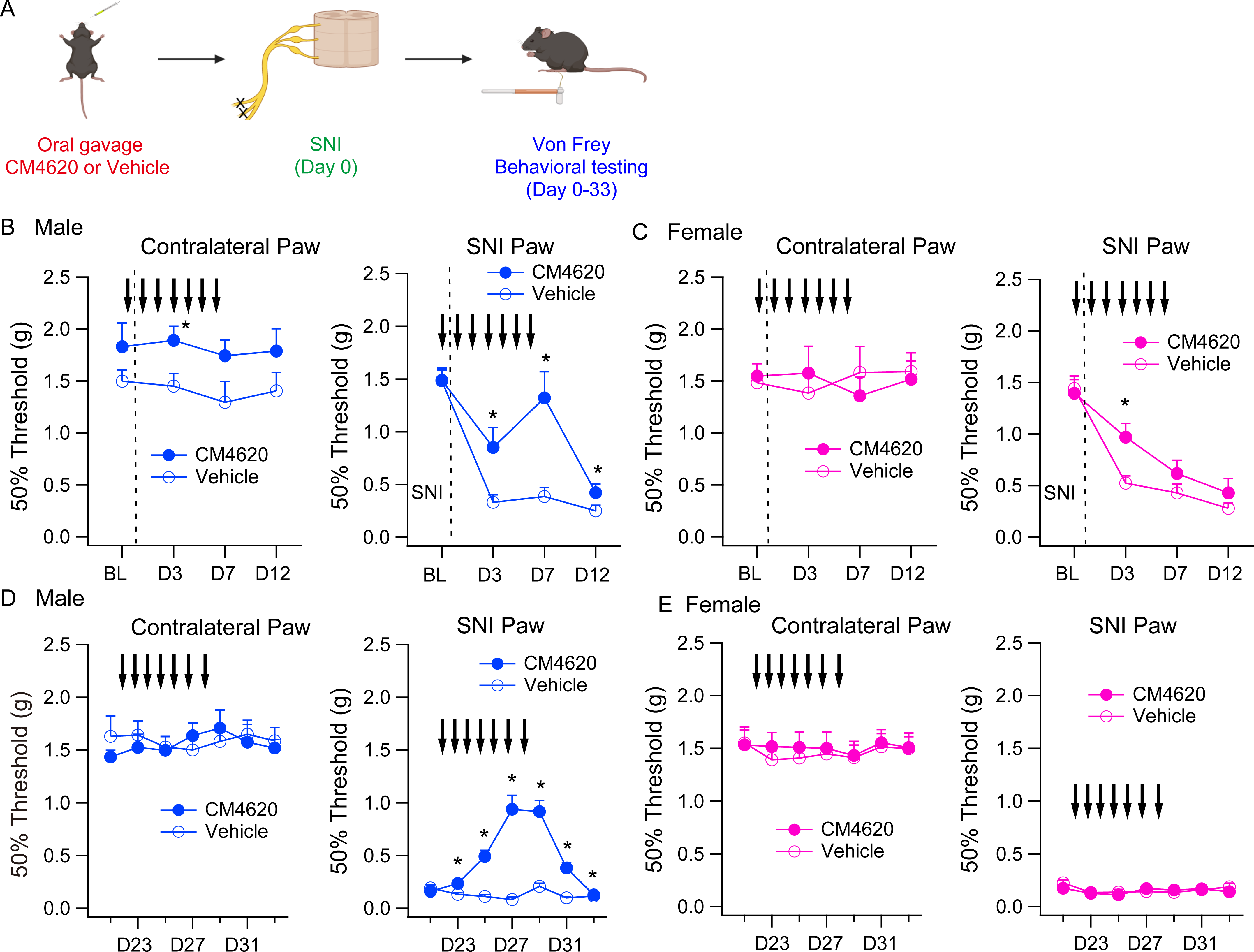
The Orai1 antagonist, CM4620, mitigates SNI-induced mechanical hypersensitivity. **(A)** A schematic of the experiment. 8-10 week old WT (Orai1*^fl/fl^*) mice were subjected to SNI (on the day indicated by the dotted line) and CM4620 (10 mg/kg) was delivered by oral gavage for 7 days as indicated by the arrows. von Frey thresholds were determined eat periodic intervals for 12 days after SNI. The mice were then rested for 9 days and retested beginning day 21. **(B,C)** von Frey thresholds in male *(B)* or female *(C)* mice before and after SNI surgery in WT (Orai1 f/fl mice) administered with either CM4620 or just the vehicle. The dotted line indicates the day of SNI surgery. **(D,E)** On day 21 following the SNI procedure, new baseline values were collected from the same mice and CM4620 or vehicle control was delivered for 7 days from the next day as indicated. N=8 mice (CM4620 group), n=7 mice (vehicle group). *: p<0.05 by unpaired T-test for comparison between CM4620 and vehicle groups.

If the effect of CM4620 is mediated by inhibition of Orai1 in microglia then one predicts that this effect would be occluded in mgOrai1 KO mice. To test this prediction we administered CM4620 in male mgOrai1KO mice to determine if delivery of the drug elicits additional analgesia compared to the protection seen in mgOrai1 KO mice alone **(Fig. S12)**. Over a duration of 31 days, these experiments indicated that neither during first few days after nerve injury nor starting 21 days after the injury, was there an effect of CM4620 in the mgOrai1 KO mice **(Fig. S12)**. One time point (day 7) following SNI surgery did show enhanced sensitivity in the mice getting CM4620, however, there was no further effect in the remaining time points. We interpret the lack of added protection in the mice getting the drug as indicative that the analgesic effect of CM4620 is likely directly due to inhibition of Orai1 in pain pathways mediated by microglia rather than by other cell types.

Together, these results indicate that *in vivo* delivery of an Orai1 antagonist mitigates mechanical hypersensitivity in the SNI model of neuropathic pain, confirming the genetic validation of Orai1 as a target for neuropathic pain (Fig. 3). Moreover, the sex-specific suppression of allodynia in male but not female mice getting the drug matches the phenotype of female microglial Orai1 KO mice, suggesting that CM4620 inhibition of microglial function may underlie the sex-specific amelioration of neuropathic pain in the wildtype mice. The ability of CM4620 to reverse hypersensitivity 3 weeks after the induction of SNI indicates that Orai1 blockers may hold promise for the development of a new line of therapeutics against neuropathic pain.

## DISCUSSION

There is strong interest in understanding the microglial checkpoints regulating neuroinflammation and several signaling cascades involving the P2X4 and P2Y receptors and p38 MAP kinase have received much attention in activating spinal microglia following nerve injury (*5*, *29*). Orai1 channels are a well-known molecular checkpoint for inflammatory processes in many immune cells (*37*, *84*, *85*), but their role in microglial-mediated neuroinflammation that underlies neuropathic pain has not been explored. In this study we show that Orai1 channels are a major mechanism for agonist-evoked Ca^2+^ signals in microglia and their opening stimulates the synthesis and release of key inflammatory cytokines. Selective deletion of Orai1 in microglia attenuated microglial activation, the production of proinflammatory spinal cytokines, and consequently, the maladaptive potentiation of excitatory synaptic transmission in the spinal dorsal horn. Surprisingly, these effects are seen only in male mice, female microglial Orai1 KO mice show no differences in these cellular and physiological endpoints. Further, behavioral analysis revealed that male, but not female microglial Orai1 KO mice show partial protection against mechanical hypersensitivity following nerve injury, mirroring the cellular phenotypes. Together, these results identify Orai1 channels as critical mediators of microglia-mediated neuroinflammation and the sex-specific effects of microglia for neuropathic pain.

Ca^2+^ imaging experiments indicated that Orai1 for mediating SOCE in spinal microglia **(Fig. 1)**, indicating that Orai1 is an essential component of the Ca^2+^ signaling toolkit in microglia (*36*, *39*, *86*). Further, the deletion of Orai1 abrogated Ca^2+^ influx arising from stimulation of microglial ATP receptors including the P2Y6 and P2Y12 receptors. This is significant as ATP released from hyper-excited or damaged neurons in the spinal cord, signaling via P2X and P2Y receptors on microglia, is considered to be a major driver of neuroinflammation and the ensuing induction of neuropathic pain (*7*, *8*, *78*, *87*). Consistent with this hypothesis, behavioral phenotyping revealed that male, but female Orai1*^fl/fl^ ^Cx3CR1-Cre/ERT2^* mice are partially protected against the induction of mechanical allodynia following SNI. The protection in male mice was robust and sustained, lasting > two weeks following nerve injury. This pattern was replicated by CM4620, a small molecule Orai1 antagonist that is currently in clinical trials for mitigating inflammation in human patients with acute pancreatitis or COVID-19 pneumonia (*47*, *48*). Importantly, CM4620 lost its protective effects in microglial Orai1 KO mice suggesting convergence of the drug’s effects on the same cellular pain pathways mediated by microglia. Moreover, cessation of CM4620 resulted in re-emergence of mechanical hypersensitivity in male mice suggesting that acute blockade of Orai1 function does not prevent the induction of neuropathic pain, but rather only its continued maintenance in male mice.

The selective effects of Orai1 deletion on spinal cytokines in male but not female mice is reminiscent of past observations that broad spectrum inhibitors of glial activation by tetracycline antibiotics, or ablation of microglia with toxins, or inhibition of microglial TLR4 or p38 signaling only protect male but not female mice in models of neuropathic pain (*22*, *23*, *26*, *27*, *51*). In addition to illustrating another striking example of a molecular perturbation yielding markedly distinct pain phenotypes in males and females, our results also bear on two key questions that are of strong interest. First, our results suggest that the sex difference in neuropathic pain in male and female mgOrai1 KO mice is directly connected to differences in proinflammatory cytokines. Male mgOrai1 KO mice exhibited lower levels of IL-6, TNF⍺, IL1-β, and BDNF in the spinal cord compared to WT counterparts following SNI **(Fig. 4)**. In female mice, by contrast, potentiation of these inflammatory mediators was unaffected by the deletion of Orai1 in microglia indicating that the behavioral sex difference is well correlated with cytokine differences. Second, our results indicate that male and female mice with differential pain regulation show differences in synaptic potentiation following nerve injury. In response to conditional deletion of Orai1 in microglia, we observed that the maladaptive potentiation of excitatory synaptic transmission following SNI is partially occluded in male but not female mice **(Fig. 4)**. Most interestingly, the pathological potentiation of excitatory synaptic transmission is correlated with the changes in proinflammatory spinal cytokines following nerve injury. Thus, these results reveal that the sex differences in pain hypersensitivity following nerve injury closely mirror changes in pathological synaptic potentiation and inflammatory cytokines in the spinal cord.

Two lines of evidence indicate that the lack of change of the upregulation of cytokine levels, excitatory synaptic enhancement, and allodynia in female Orai1*^fl/fl^ ^Cx3CR1-Cre/ERT2^* mice following SNI is not because Orai1 is non-essential for the physiology of female microglia. First, deletion of Orai1 in the isolated female microglia abrogated SOCE equally from isolated microglia in both males and females **(Fig. S2)**. This finding indicates that Orai1 is indeed essential for agonist-evoked SOCE in both males and females. Second, deletion of Orai1 impaired cytokine synthesis (IL-6, TNF⍺, MCP1, and PGE_2_) from primary cultures of microglia in both males and females **(Fig. 6)**, indicating that the difference between males and females in the cytokine synthesis is not a cell autonomous feature of microglia.

What then could be the basis of the *in vivo* male-female difference in the microglial Orai1 KO mice? Several scenarios are possible. We observed that there was no statistical difference in activation of the microglial marker IBA1 between male and female mice in the spinal cord **(Fig. 6)**, suggesting that nerve injury evokes similar degree of microglia proliferation in both sexes in line with previous studies (*23*). However, ablating expression of Orai1 did not dampen the degree of microgliosis as assessed by IBA1 in female mice, in contrast to male mice where significant reduction in IBA1 was readily detected **(Fig. 6)**. This key result indicates that *in vivo*, SNI activates microglia in female mice with equal efficacy even in the absence of Orai1 to override Orai1 contributions to microgliosis. How might this occur? Recent evidence indicates that female mice may show greater dependence on T cells compared to male mice, and are protected against neuropathic pain primarily when T-cells and microglia are both depleted (*23*). Microglia and T cells can interact in numerous ways in the CNS including cell-to-cell contact and cytokine mediated communication to promote neuroinflammation (*88*). We postulate that in female rodents lacking Orai1, microglial activation is mediated in large part by stronger inputs (relative to male mice lacking Orai1) from T cells. Because these T-cell derived inputs are not impaired, microglia in female Orai1 KO mice may maintain strong levels of microglial reactivity to override the loss of Orai1. An alternative possibility is that Orai1 may not be engaged following SNI in female microglia, as evidenced by IBA1 labelling which is unaffected in Orai1 cKO mice relative to WT controls. This may occur because the upstream receptor (e.g., P2X4) that turns on IBA1 expression may differ between males and females resulting in differential regulation upon Orai1 deletion. These possibilities are of course not mutually exclusive but the results presented here provide a framework for evaluating these and other hypotheses.

## MATERIALS AND METHODS

### Transgenic mice

Male and female C57BL/6 mice were used in this study and cared for in accordance with institutional guidelines and the *Guide for the Care and Use of Laboratory Animals*. All animals were group-housed in a sterile ventilated facility, under standard housing conditions (12:12 hour light/dark cycle with lights on at 7:00 a.m. and temperatures of 20 to 22°C with ad libitum access to water and food), and maintained with in-house breeding colonies. Male and female mice were used in approximately equal numbers. All research protocols were approved by the Northwestern University Institutional Animal Care and Use Committee. The Orai1 *^fl/fl^* mice have been described previously (*46*). Tissue-specific deletion of Orai1 in microglia was accomplished by crossing Orai1*^fl/fl^* mice with Cx3CR1-Cre/ERT2 mice (Jax labs) to generate Orai1*^fl/fl^ ^Cx3CR1-Cre/ERT2^* mice. Cre recombinase activity was induced as previously described (*89*). Briefly, 4-5-week old mice were treated with 8 mg tamoxifen (TAM, T5648, Sigma-Aldrich, USA) solved in 200 ul corn oil (C8267, Sigma) injected intraperitoneally at two time points 48 h apart. *In vitro* recombination was achieved using OH-TAM (H7904, Sigma-Aldrich, USA) at a final concentration of 1 µM to 25 cm^2^ glia culture flasks at least 3 days before harvesting microglia. Ethanol was used as a solvent control for in vitro experiments.

### Primary cultures

Primary microglia were isolated from neonatal (P0 to P3) mice from astrocyte-glial cultures which were grown as previously described for hippocampal tissue (*42*). Briefly, spinal cord was dissected, and meninges were removed under a dissection microscope in 4℃ dissection medium [10 mM HEPES in Hanks’ balanced salt solution (HBSS)]. After enzymatic dissociation with 0.25% Trypsin (Invitrogen) and DNase (Roche, Mannheim, USA) for 15 min in a 37℃ water bath, tissue was washed twice with HBSS and dissociated gently by trituration in culture media consisting of Dulbecco’s modified Eagle’s medium with 10% fetal bovine serum and 1% penicillin-streptomycin solution. Dissociated cells were filtered through a 70-um strainer to collect cell suspension and cultured in 25-mm^2^ tissue culture flasks on poly-D-lysine-coated (Sigma-Aldrich, Schnelldorf, USA) with 10 ml of medium. When cultures were made for Ca^2+^ imaging experiments to evaluate SOCE differences in males and females, cells were cultured separately from each pup. The spinal cord was dissected, cells isolated as described above, and cultured in T25 flask for two weeks. For ELISA experiments evaluating cytokine production in cultured microglia (Fig. 6), 2 spinal cords from neonatal mice of the same sex were used per flask. Half of the medium was exchanged every 3 to 4 days until cells reached near confluence (7 to 10 days in vitro). Microglia were collected by forcefully shaking by hand for 1 min., or by shaking cells in an incubator for 1 hour at 200 rpm, and centrifuged for 5 min. The supernatant was removed, and the cell pellet was resuspended in culture medium. Cells were plated on poly-D-lysine-coated glass-bottom dishes (MatTek, 14-mm diameter, 10,000 to 15,000 cells per coverslip) or 24-well plates (15,000 to 20,000 cells per well). Plated microglia were maintained in the incubator and used in experiments after 1 week for the Ca^2+^ imaging and ELISA studies shown in Figures 1 and 6 and 2 weeks for the experiments shown in Supplementary Figure 2, which used microglia cultured separately from 1 pup grown individually. Half of the medium was exchanged with fresh medium every 4 days.

### Isolation of microglia from adult mice

Protocol for isolation of adult microglia in mice were performed as previously described (*90*). Briefly, Twelve-week-old mice were anaesthetized with isoflurane and perfused intracardially with cold PBS. Brain and spinal cords were extracted from each mouse and minced in DMEM/F12 with 1% Pen-Strep and 1% glucose. Minced tissue was then digested in DMEM/F12 supplemented with DNAseI (20U/ml, Roche Diagnostics, 10104159001), papain (1mg/ml, Worthington Biochemical Corporation, LK003178), and dispase II (1.2U/ml, Millipore Sigma, D4693-1G) at 37° for 20 minutes. Afterward, the dissociation was neutralized with DMEM/F12 with 10% heat inactivated FBS and 1% pen-strep. Tissue was washed twice with DMEM/F12 supplemented with 1% glucose and 1% pen-strep. Tissue was mechanically dissociated by pipetting up and down with P1000. Once the large clumps had settled, the upper part was collected and transferred to a new tube. This was repeated once more with a P1000 with fresh media, and then repeated one more time with P200. The collected cells from the dissociation were filtered through a 70µm cell strainer and centrifuged. The supernatant was removed, and the cell pellet was resuspended in 37% stock isotonic percoll (SIP, 3.7ml of SIP solution and 5.3 ml of 1xHBSS). SIP solution was 1 part 10xHBSS (Gibco, 14185-052) and 9 parts percoll (Sigma, P493). The 70% SIP layer was slowly added below the 37% SIP cell containing layer, and the 30% SIP solution was added above the 37% SIP layer. Finally, 2mL of 1XHBSS was added above the 30% SIP layer and centrifuged for 40 minutes at 300xg and 18°C with no brake. After centrifugation, the 2 mL of solution was collected at the 70-37% interphase and transferred to a clean tube. The interphase was washed with 6mL of 1XHBSS and centrifuged at 500xg for 7 minutes. Cell pellet is then resuspended in 500µl of HBSS and transferred to 1.5mL tube followed by washing 3 times with 1xHBSS. Cells were then used for RNA extraction and quantitative RT-PCR.

### Ca^2+^ imaging

Primary spinal microglia grown on poly D-lysine coated glass-bottom dishes were loaded with Fura-2 by incubating cells in 2 uM Fura-2-AM (Invitrogen) in growth medium for 30 min at 37℃. Fura-2-containing medium was washed off, and cells were incubated for an additional 10 min before imaging. All experiments were performed at room temperature. Single cell [Ca^2+^]_i_ measurements were performed as described previously (Toth et al., 2019). Image acquisition and analysis were performed using SlideBook (Denver, CO). Dishes were mounted on the stage on Olympus IX71 inverted microscope, and images were acquired every 6 sec at excitation wavelengths of 340 and 380 nm, and an emission wavelength of 510 nm. For data analysis, regions of interest were drawn around single cells, background was subtracted, and *F_340_/F_380_* ratios were calculated for each time point. A rise in the ratio of emission when excited at 340 nm over the ratio when excited at 380 nm indicated a rise in [Ca^2+^]_i_.

[Ca^2+^]_i_ was estimated from *F_340_/F_380_* ratio using the standard equation: [Ca^2+^]_i_ = βK_d_ (R-R_min_)/(R_max_-R), where R is the *F_340_/F_380_* fluorescence ratio and values of R_min_ and R_max_ were determined from an in vitro calibration of Fura-2 pentapotassium salt. β was determined from the *F_340_/F_380_* ratio at 380 nm and K_d_ is the apparent dissociation constant of Fura-2 binding to Ca^2+^ (135 nM). For each cell, the rate of SOCE (Δ[Ca^2+^]_i_/Δt) was calculated from the slope of a time fitted to three points (18 sec) after the re-addition of 2 mM Ca^2+^ Ringer’s solution. Baseline [Ca^2+^]_i_ was calculated by averaging [Ca^2+^]_i_ values over 2-min baseline for each experiment. Store release was calculated by measuring the area under the curve during TG application in Ca^2+^-free solution. The standard Ringer’s solution used for these experiments contained the following (in mM): 155 NaCl, 4.5 KCl, 10 D-glucose, 5 HEPES, 1 MgCl_2_, 2 CaCl_2_. The Ca^2+^-free Ringer’s solution contained 3 mM MgCl_2_, 1 mM EGTA (Sigma-Aldrich) and no added CaCl_2_. pH was adjusted to 7.4 with 1 N NaOH. Stock solution of thapsigargin (TG) was dissolved in DMSO and used at the indicated concentration.

### Immunostaining of cultured microglia

Isolated microglia were seeded on poly-d lysine coated glass coverslips and fixed at room temperature using 4% PFA for 15 minutes. Cells were then washed twice with PBS, permeabilized and blocked using 4% BSA/4%NGS/0.1% TritonX-100 for 1hr. Cells were then washed twice using PBS and incubated overnight with 1:1000 Iba1 (Abcam) in PBS/4%BSA at 4℃. Next day cells were washed with PBS containing 0.1% Tween- 20 and then incubated by secondary goat rabbit Alexa Fluor 488 antibody (Thermofisher, 1:1000 in PBS/4%BSA) at room temperature in the dark. Cells were then washed 3times using PBS/0.1% Tween and were mounted using ProLong Gold Antifade Mount medium with DAPI (P36931) and imaged on an upright Nikon A1R confocal microscope.

### Quantitative RT-PCR

Microglia cultures from mouse pup spinal cords were lysed for total RNA extractions using RNeasy Plus Mini Kit (74134, Qiagen). cDNA generation was performed using High-Capacity cDNA Reverse Transcription Kit (4368814, Applied Biosystems by Thermo Fisher Scientific). PowerUp SYBR Green Master mix (A25741, Applied Biosystems by Thermo Fisher Scientific) was used for quantitative PCR (qPCR) and followed according to manufacturer’s instructions. For qPCR, each well contained 7 ng of cDNA and final primer concentration of 500 nM. The qPCR was run using CFX Connect Real-Time System from Bio-Rad, and data was analyzed using Bio-Rad CFX Maestro software. Sequences of primers: Orai1 forward, 5’-AGACTGCCTGATCGGATGGC-3’, Orai1 reverse, 5’-TTGTCCCCGAGCCATTTCCT-3’; Gapdh forward, 5’-AGGTCGGTGTGAACGGATTTG-3’, Gapdh reverse, 5’-ATGTA GACCATGTAGTTGAGGTCA-3’;18S forward, 5’-TGCGAGTACTCAACACCA ACA-3’, 18S reverse, 5’-CTGCTTTCCTCAACACCACA-3’.

### ELISA analysis

For analysis of cytokine levels in spinal tissue *in vivo*, the spinal cord (5 mm in length) at the lesion epicenter was dissected out from each animal (5 animals per group) 7 days after SNI surgery. Spinal cord tissue was homogenized by mechanical trituration followed by sonication in radio-immunoprecipitation assay (RIPA) lysis buffer (1 ml; Santa Cruz Biotechnology, Inc., Santa Cruz, CA, USA) and the cytokine concentration were measured using mouse ELISA kits (Invitrogen Life Technologies, Carlsbad, CA, USA) according to the manufacturer’s instructions. Cytokine levels were expressed in pg/mg.

For the *in vitro* cytokine measurements from cultured cells, primary microglia isolated from spinal cords were seeded into 24-well plates at a density of 20,000 cells per well. Serum was removed from the media 24 hours prior to treatments being added to the wells. Cells were treated with DMSO, LPS (1 ng/ml), or TG (0.25 µM) and phorbol 12,13-dibutyrate (25 nM) for 18 hours. Supernatants were subsequently analyzed using mouse ELISA kits for the release of TNFα (R&D Systems, MTA00B), IL-6 (RayBiotech, ELM-IL6-1), PGE_2_ (Cayman Chemical, 514010), and MCP-1 (RayBiotech, ELM-MCP1-CL-5). All kits were used according to the manufacturers’ instructions. Release of pro-inflammatory factors is expressed in concentration of pg/ml.

### Preparation of spinal cord slices

Spinal cord slices obtained from mice at 7 days after SNI surgery were used for electrophysiological assessments. The mice were deeply anesthetized with isoflurane and then subjected to a rapid intra-cardiac infusion of ice-cold oxygenated high-sucrose artificial cerebrospinal fluid (sACSF) containing (in mM): 95 NaCl, 1.8 KCl, 1.2 KH_2_PO_4_, 0.7 CaCl_2_, 1 MgCl_2_, 26 NaHCO_3_, 50 sucrose, and 15 glucose. The lumber segments (L3-L5) were quickly removed and immersed in ice-cold oxygenated sACSF. The pH was adjusted to pH 7.4 (osmolality 300-310 mOsm), and the fluid was oxygenated with 95% O_2_ and 5% CO_2_. To identify the ipsilateral side, a sharp knife was used to make a deep mark in the contralateral side. The spinal cord was then transversely sliced into 400 um sections using a tissue slicer (Compresstome model VF-200-0Z, Precisionary Instruments). Slices were transferred to a recovery ACSF chamber containing (in mM): 125 NaCl, 2.4 KCl, 1.2 Na_2_PO_4_, 1 CaCl_2_, 2 MgCl_2_, 25 NaHCO_3_, and 25 glucose, while the slices were maintained at 30 ℃. Individual slices were transferred to a recording chamber and perfused with normal ACSF containing 2 mM CaCl_2_.

### Slice electrophysiology

The lamina II (substantia gelatinos area) in the superficial dorsal horn was visually identified as translucent wide band of tissue in spinal slices under differential interference contrast imaging. EPSCs and IPSCs were recorded from lamina II neurons using the whole-cell patch-clamp technique with an Axopatch 200B amplifier interfaced to an ITC-18 input-output board and iMac G5 computer. Currents were filtered at 2 kHz with a four-pole Bessel filter and sampled at 5 kHz. Data acquisition and analysis were performed using in-house routines developed on the Igor Pro platform. Voltage-clamped recordings were performed in the whole-cell recording mode with a pipette potential of -70 mV. Patch electrodes were pulled from borosilicate capillary glass using a vertical pipette puller and had a resistance of 4-6 MΩ. The pipettes were filled with an intracellular solution containing (in mM): 135 K-gluconate, 5 KCl, 5 ethylene glycol tetraacetic acid (EGTA), 5 Mg-ATP, 5 HEPES, 0.5 CaCl_2_ and 2 MgCl_2_ (adjusted to pH 7.2 with Tris base) for recording of EPSCs or 95 CsF, 25 CsCl, 10 HEPES, 10 EGTA, 2 Mg-ATP, 0.3 Na_3_-guanosine triphosphate, 10 QX-314, 5 5 tetraethylammonium chloride and 5 4-aminopyridine (pH 7.3 with KOH or CsOH) for IPSCs. Spontaneous EPSCs were recorded in the presence of picrotoxin (50 µM) and D-(-)-2-Amino-5-phosphonopentanoic acid (D-APV) (50 µM). Spontaneous IPSCs (sIPSCs) were isolated by the administering 6-cyano-7-nitroquinoxaline-2,3-dione (CNQX) (10 µM) and D-APV (50 µM) to the slices in the extracellular solutions. Miniature EPSCs (mEPSCs) and IPSCs (mIPSCs) were recorded in the presence of tetrodotoxin (TTX) (1 µM).

### Immunohistochemistry of spinal cord slices

Mice were anaesthetized with isoflurane and perfused intracardially with cold PBS followed by 10% formalin 14 days after SNI. Spinal cords were extracted, and the lumbar area was isolated and fixed in 10% formalin solution overnight. The next day spinal cords were placed in 30% sucrose PBS solution. Spinal cords were sectioned at 40µm thickness. Sections were washed and blocked with 10% goat serum in TBS-T for two hours at room temperature. Slices were incubated with rabbit anti-IBA1 primary antibody (1:1000, Wako Chemicals USA, 019-19741) at 4°C overnight. Sections were washed and incubated with secondary antibody (1:1000, goat anti-rabbit Alexa594) and dapi for two hours at room temperature. Sections were washed and mounted with VectaShield (Vector Labs, H-1200-10). Images of the spinal cord and spinal dorsal horns were acquired using a Nikon A1R confocal microscope with a 25x objective. The images of whole spinal cord were taken by stitching images together and used to identify L4-L5 area of the spinal cord. If a section had IBA1+ staining in the ventral horn, a z-stack image was acquired of the dorsal horn. All image acquisition settings were maintained between sexes, genotypes, and injury groups. The number of stacked images and step size was consistent between sexes, genotypes, and injury groups. Prior to quantification, the z-stack images were flattened to maximum intensity projections, and then subsequently processed for percent area of IBA+ staining. Background was subtracted and the dorsal horn was outlined to create a region of interest (ROI). The ROI was used to quantify the percent area of IBA1+ staining detectable above the background. Images were analyzed with NIS Elements software and analysis parameters were kept the same between sexes, genotypes, and injury groups.

### SNI surgery

SNI surgeries were performed following the procedures described previously (*61*). Under anesthesia with isoflurane in O_2_ (induction: 3% isoflurane; maintenance: 2% isoflurane), the left sciatic nerve was exposed after sectioning the skin of the posterior aspect of the thigh and incision through the fat pad covering the popliteal fossa. The common peroneal and the tibial nerves were identified and tightly ligated with 5-0 silk thread and transected distal to the ligation, removing 2-4 mm length of each nerve. The sural nerve remained intact and any contact with or stretching of this nerve was carefully avoided. Muscle and skin were closed in two distinct layers. The sciatic nerve trifurcation was only exposed without ligation and transection for sham surgeries.

### Mechanical withdrawal threshold assessment

The 50 % paw withdrawal threshold (PWT) of both hind paws was assessed using von Frey filaments as previously described (*91*). Briefly, the animals were placed individually under separate transparent Plexiglass chambers positioned on a wire mesh floor. Approximately 30 min were allowed for habituation. The mice were sequentially applied a series of von Frey filaments (0.02-4.0 g; Stoelting CO, USA) to the lateral plantar surface of the hind paw. Filaments were perpendicularly applied to the lateral plantar surface of the hind paw in either ascending or descending strengths to determine the filament strength closest to the withdrawal threshold. Each filament was applied for a maximum of 2 s at each interval. Brisk withdrawal and hind paw licking were considered positive as nociceptive responses. Given the response pattern and the force of the final filament, 50% response threshold was calculated. Other behavioral changes were also recorded. All tests were conducted between 1 and 4 PM by an examiner who was blind to the experiments.

### Drug administration

Stocks of BTP (catalog#203890; Sigma-Aldrich) and the powdered form of CM4620 (Calcimedica) were dissolved in DMSO and applied at dilutions of 1000-fold. For the behavioral studies in Figure 7, 40 mg of CM4620 was mixed with and 9.6 ml of vehicle (0.5% methyl cellulose and 1% Tween 80) for 10 mg/kg (assuming 10 ml/kg and 25% loading of CMEX-128). All treatments were delivered daily to the mice via oral gavage 7 days from the day of surgery as indicated in the figure legend. CNS penetration of CM4620 was confirmed by determining the ratio of compound concentrations (measured by LC-MS/MS) in brain tissue extracts to those in plasma after oral or IV administration. Under these conditions, the brain to plasma ratio ranged from 2.0 to 2.6 in both rats and mice and did not accumulate with repeated dosing. For the drug re-administration studies in Figure 7D,E beginning day 22 post injury, CM4620 or vehicle was administered to mice for 7 days.

### Data analysis

All data are expressed as means ± SEM. Statistical analysis was calculated using SigmaPlot version 13.0 (Systat Software) with a confidence level of 95%, and results with P < 0.05 were considered statistically significant (*p<0.05, **p<0.01, ***p<0.001). For datasets with two groups, statistical analysis was performed with a two-tailed Student unpaired *t* test to compare between groups. For datasets greater than two groups, one-way ANOVA followed by Tukey post-hoc test was performed to compared between groups. Miniature EPSCs (mEPSCs) and IPSCs (mIPSCs) were analyzed using MiniAnalysis software (Synaptosoft Inc., GA, USA). The threshold for detection of events were set at 5 pA (for EPSCs) or 10 pA (for IPSCs).

## ACKNOWLEDGEMENTS

We thank members of the Prakriya laboratory for helpful discussions and the behavioral core of the Northwestern University Pain Center for assistance with the von Frey analysis. This work was supported by NIH grants R01NS057499 and R21NS122347 to MP, P50 DA044121 to AVA, and CIHR grant FDN-154336 to MWS.

## AUTHOR CONTRIBUTIONS

ST carried out the Ca^2+^ imaging, electrophysiology, *in situ* cytokine measurements, and behavioral analysis, KED performed Ca^2+^ imaging, *in vitro* cytokine analysis, immunohistochemistry of spinal cord sections, and behavioral analysis, MC, MEM, and MRT performed the SNI surgeries and assisted with the behavioral analysis, MM and MY devised the cell isolation and spinal cord dissection procedures and performed Ca^2+^ imaging, and SG and MWS conceived and carried out the RNA-seq analysis. ST, KED, MWS, KAS, AVA, and MP conceived and designed experiments. ST, KED, KAS, MWS, and MP wrote and edited the paper.

## DATA AND MATERIALS AVAILABILITY

All data needed to evaluate the conclusions in the paper are present in the paper or the Supplementary Materials and from the corresponding author upon request. CM4620 was obtained via an MTA from Calcimedica.

## SUPPLEMENTARY INFORMATION

**Figure S1.**
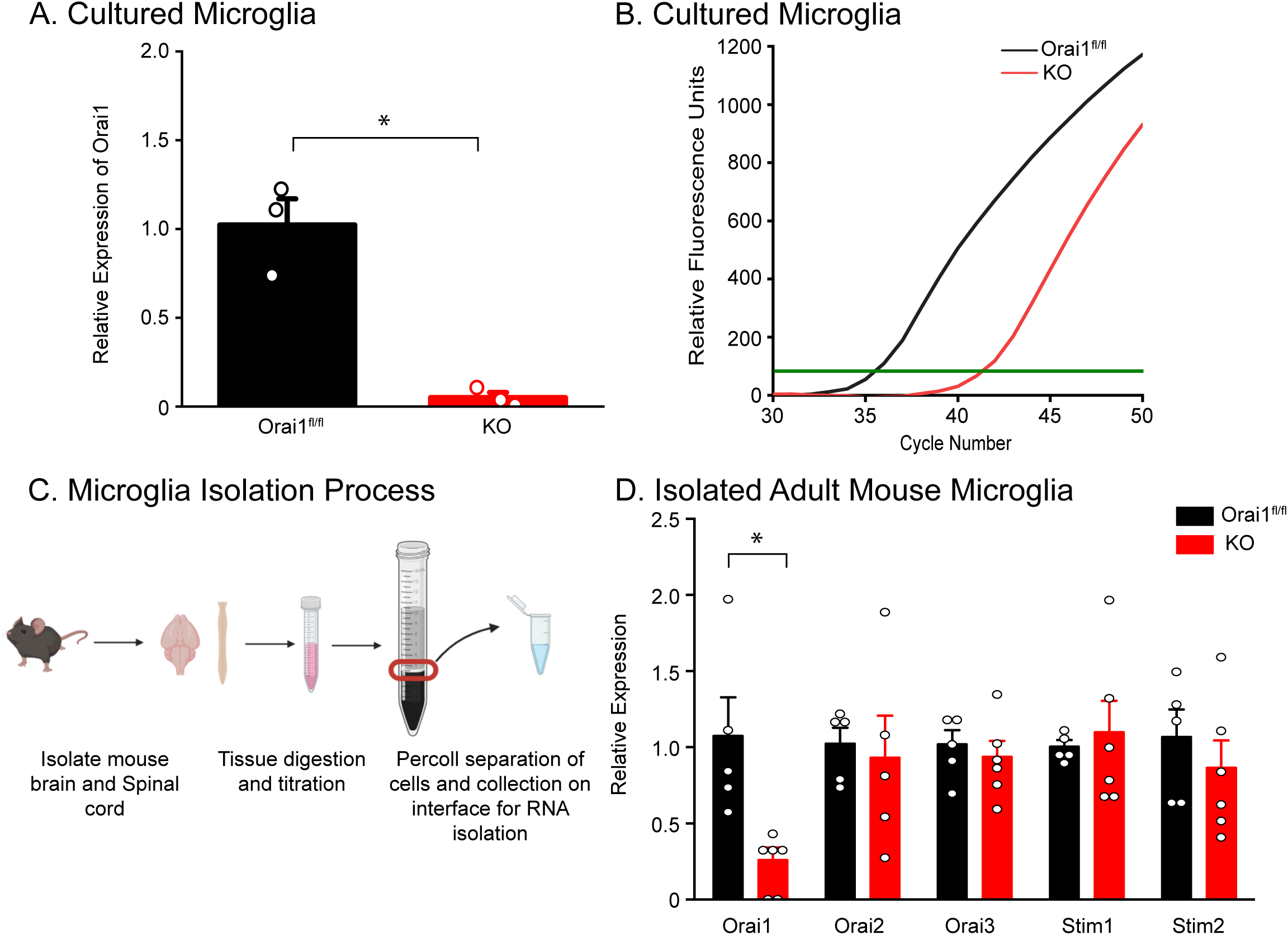
Analysis of Orai1 mRNA expression in primary microglia. Primary microglia were isolated from the spinal cord from P0-P3 pups and Orai1 mRNA was assessed by real-time qPCR. **(A)** Relative expression of Orai1 mRNA in WT (Orai1 *^fl/fl^*) and Orai1 KO (Orai1 *^fl/fl^ ^CX3CR1-Cre/ERT2^*) microglia exposed to tamoxifen. mRNA was normalized to GAPDH and 18S mRNA, and the mean of the wildtype group was normalized to 100%. **(B)** Real-time monitoring of the fluorescence emission of SYBR Green I during PCR amplification of Orai1 cDNA obtained by reverse transcription from mRNA harvested from spinal microglia cultures from WT and Orai1 KO mice. The dashed line denotes noise level. N= 3. P<0.05 by unpaired student’s T-test. **(C)** Protocol for isolation of microglia from the adult brain and spinal cord. A percoll gradient centrifugation step was used to separate adult primary microglia from other cells. **(D)** RT-qPRC analysis of Orai and STIM isoforms from freshly isolated adult microglia. N=5-6. P<0.05 by unpaired student’s T-test.

**Figure S2.**
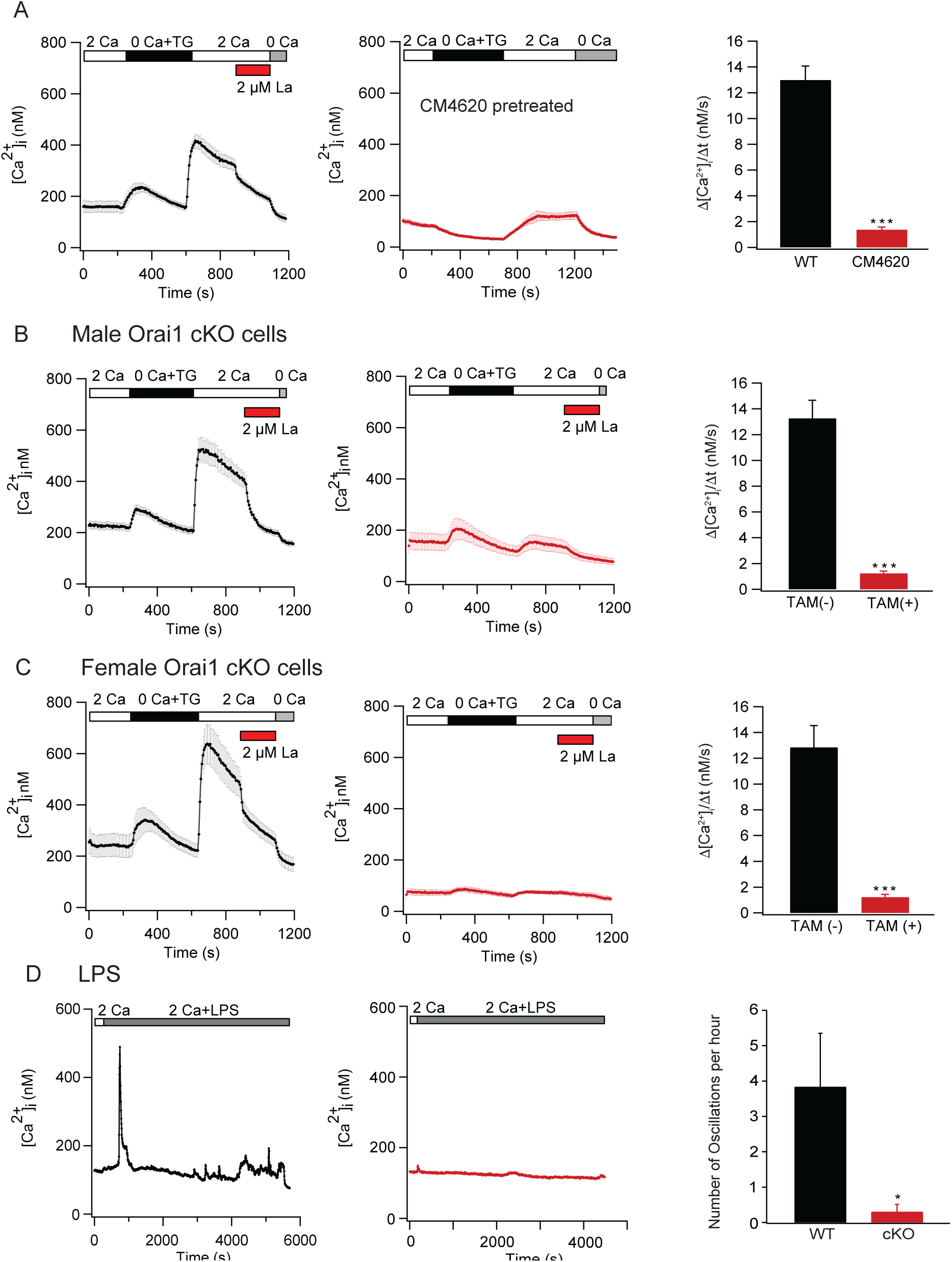
Analysis of SOCE and effects of Orai1 deletion in microglia isolated from male and female mice. **(A)** Inhibition of SOCE by the Orai1 antagonist, CM4620 in WT (Orai1*^fl/fl^*) microglia. SOCE was initiated by depleting Ca^2+^ stores with 1µM thapsigargin (TG) in Ca^2+^ -free medium followed by re-addition of extracellular Ca^2+^ at t= 500 s. The bar graph on the right is the quantification of the rates of Ca^2+^ entry following extracellular Ca^2+^ re-addition. n= 60 cells for CM4620 treated cells. **(B,C)** Analysis of SOCE in male and female microglia cultured separately from individual postnatal mice. Microglia isolated from postnatal pups were cultured separately (from each mouse) for 14 days. SOCE was initiated by depleting ER Ca^2+^ stores with TG as described above. 2 µM La^3+^ was added to the extracellular Ringer’s solution at the indicated times. The right graph summarizes the rate of Ca2+ influx following re-addition of extracellular Ca^2+^. n=84 (males, no tamoxifen) and n=96 (males, tamoxifen). n=109 (females, no tamoxifen) and n=56 (females, tamoxifen) ***: p < 0.0001 by Mann-Whitney Rank sum test. **(D)** Example traces of stimulation of WT (Orai1*^fl/fl^*) and Orai1 KO (Orai1 *^fl/fl^ ^CX3CR1-Cre/ERT2^*) microglia with LPS (1µg/ml) stimulation in the extracellular Ca^2+^ Ringer’s Solution. Ca^2+^ oscillations were measured by calculating change of the intercellular calcium for each time point after addition of LPS. The number of time points that had a change greater than or equal to 20 nM were counted and normalized per hour. n=12 cells (Orai1*^fl/fl^*) and n=10 cells (Orai1 *^fl/fl^ ^CX3CR1-Cre/ERT2^*). *: p<0.05 by unpaired student’s T-test.

**Figure. S3.**
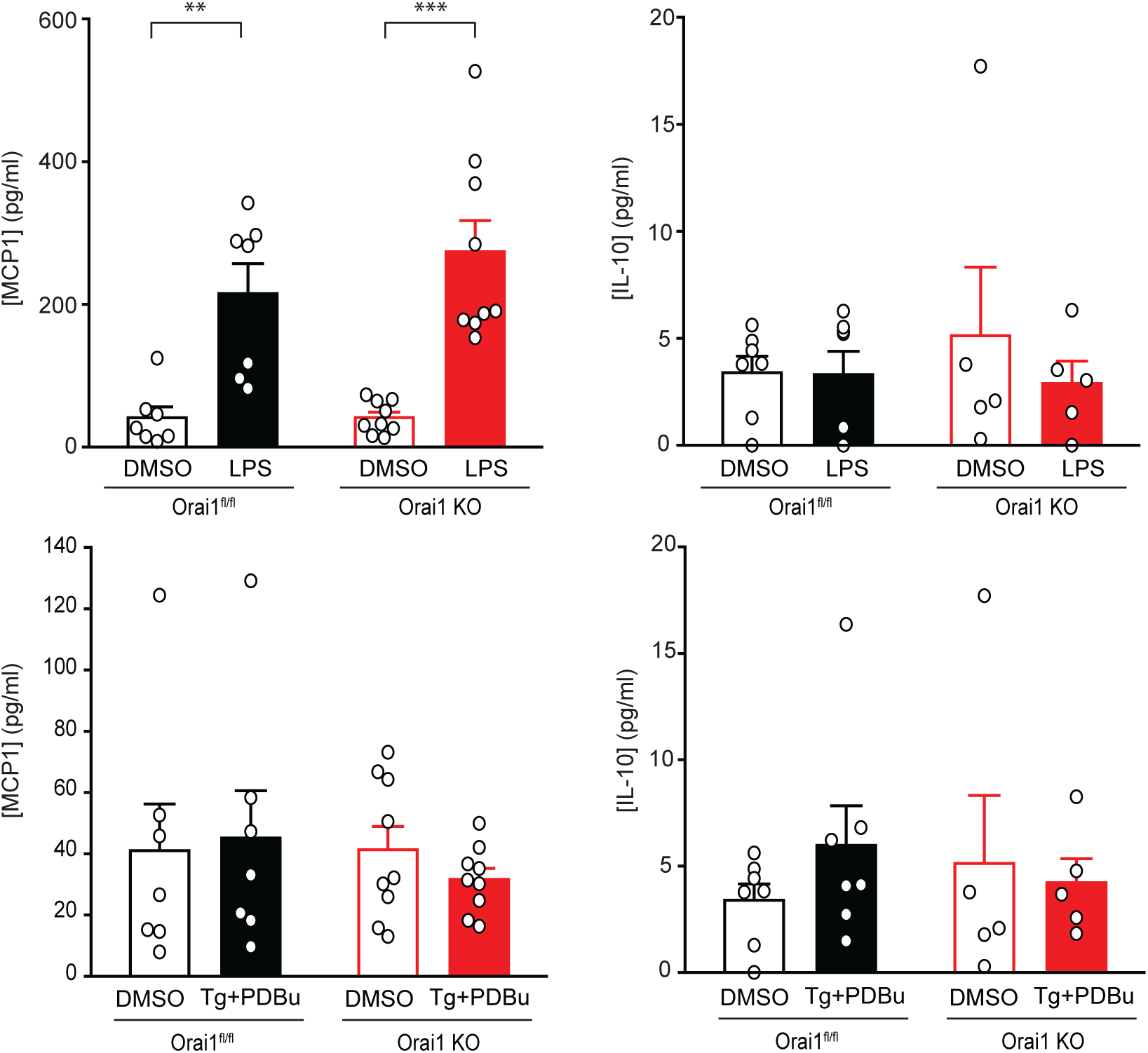
Analysis of MCP1 and IL-10 from spinal microglia. MCP1 and IL-10 levels were assessed in the supernatant 18 hours following cell stimulation via ELISA. n=5-9 measurements from 10-18 mice. **: p<0.01 and ***: p<0.001 by two-way ANOVA followed by Tukey test for comparison between multiple groups. Deletion of Orai1 in microglia does not affect induction of MCP1. Moreover, IL-10 is not affected by genotype or the treatment.

**Figure S4.**
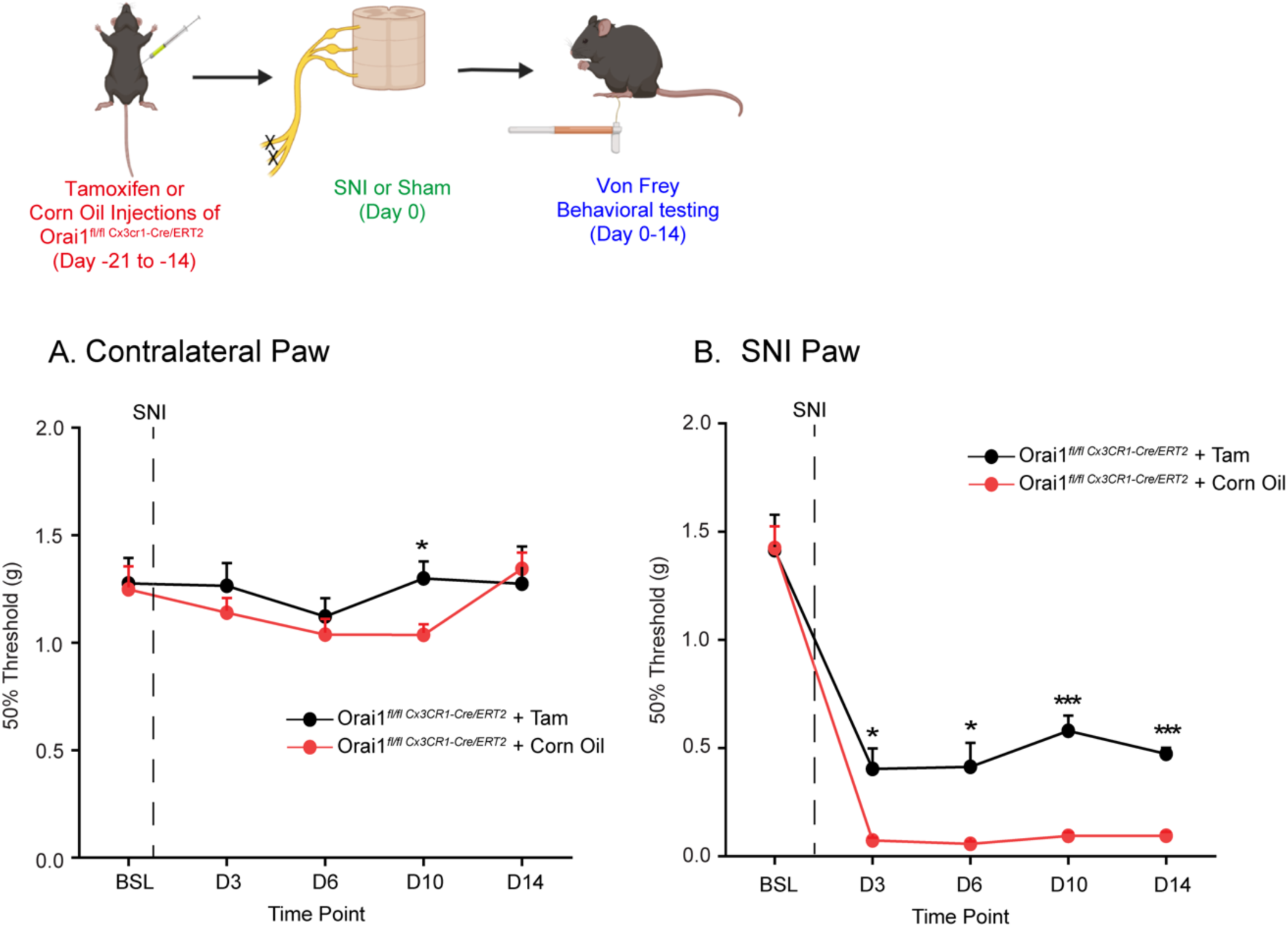
Analysis of allodynia in male Orai1*^fl/fl^ ^CX3CR1-Cre/ERT2^* mice with and without tamoxifen exposure. Mechanical sensitivity measured by von Frey thresholds in male Orai1*^fl/fl^ ^CX3CR1-Cre/ERT2^* mice injected with either tamoxifen or vehicle alone (corn oil). The dotted line indicates that day when SNI was performed. Only Orai1*^fl/fl^ ^CX3CR1-Cre/ERT2^* mice administered with tamoxifen to induce Orai1 deletion are partially protected from allodynia indicating that the protection is not due to Cre expression. n=9 (corn oil) and n=8 (tamoxifen). *: p < 0.05; ***:p <0.001 by unpaired T-test comparing male Orai1*^fl/fl^ ^CX3CR1-Cre/ERT2^* mice with and without tamoxifen administration for each time point.

**Figure S5.**
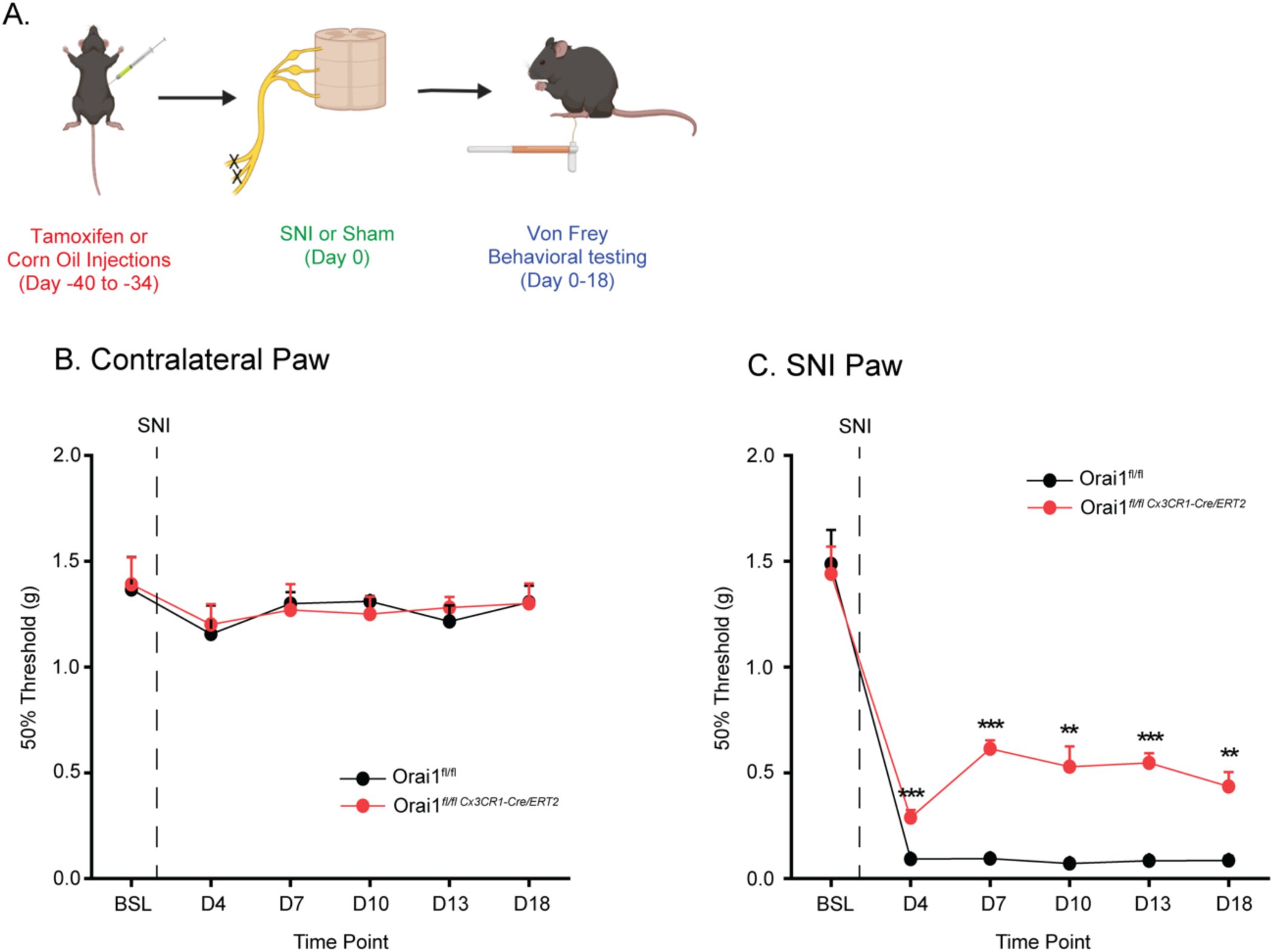
Analysis of pain hypersensitivity in male Orai1*^fl/fl^ ^CX3CR1-Cre/ERT2^* mice with increased waiting period following tamoxifen administration. Mechanical sensitivity measured by von Frey thresholds in male Orai1*^fl/fl^ ^CX3CR1-Cre/ERT2^* mice injected with either tamoxifen or vehicle alone (corn oil). The dotted line indicates that day when SNI was performed. Following the last tamoxifen injection, 34 days were allowed prior to SNI. Paw withdrawal thresholds were measured for 18 days following SNI. Orai1*^fl/fl^ ^CX3CR1-Cre/ERT2^* mice administered with tamoxifen are partially protected from allodynia throughout the measurement period even at 54 days following tamoxifen injection suggesting that the mitigation is not mediated by infiltration of peripheral monocytes expressing CX3CR1. n=6 (Orai1*^fl/fl^*; males) and n=7 (Orai1 *^fl/fl^ ^CX3CR1-Cre/ERT2^*; males). **: p < 0.01; ***:p < 0.001 by unpaired T-test comparing each time point.

**Figure S6.**
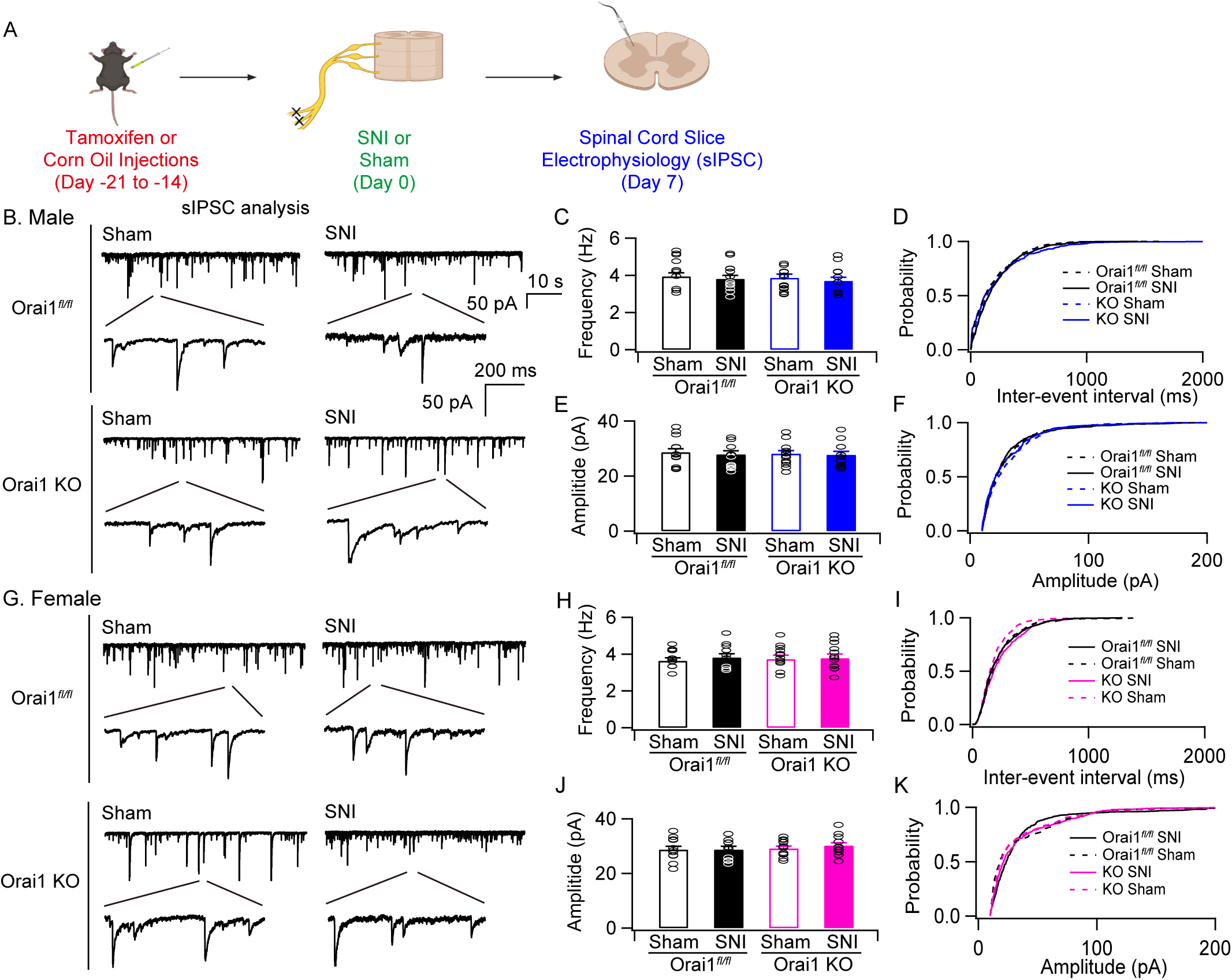
SNI does not affect the frequency or amplitude of spontaneous IPSCs. **(A)** A schematic of the experimental plan. **(B)** Examples of sIPSCs recorded from dorsal horn neurons of L4 spinal cord slices (-70 mV) in lamina II. Excitatory synaptic transmission was blocked by the AMPA and NMDA receptor blockers NBQX and D-APV. **(C)** Bar graphs (mean±sem) summarizing the sIPSC frequency. **(D)** Cumulative probability distribution of sIPSCs in the indicated groups. **(E)** Summary (mean±sem) of the amplitude of the sIPCS. **(F)** cumulative probability distribution of the sIPSC amplitudes. **(G)** Example sIPCS traces at how and high tome resolution from slices from female mice. **(H, J).** Summary of the frequency and amplitude of sIPSCs in female mice in the indicated groups. **(I,K)** cumulative probability distributions of the sIPSC frequency and amplitudes. N values are as follows: Male WT mice: n=17 cells (sham), n=17 (SNI). Male Orai1 KO mice: n=17 cells (sham), n=17 cells (SNI). Female WT mice: n=12 cells (sham), n=12 cells (SNI). Female Orai1 KO mice: n=12 cells (sham), n=12 cells (SNI). Statistical analysis using ANOVA followed by Tukey test for comparison of multiple groups.

**Figure S7.**
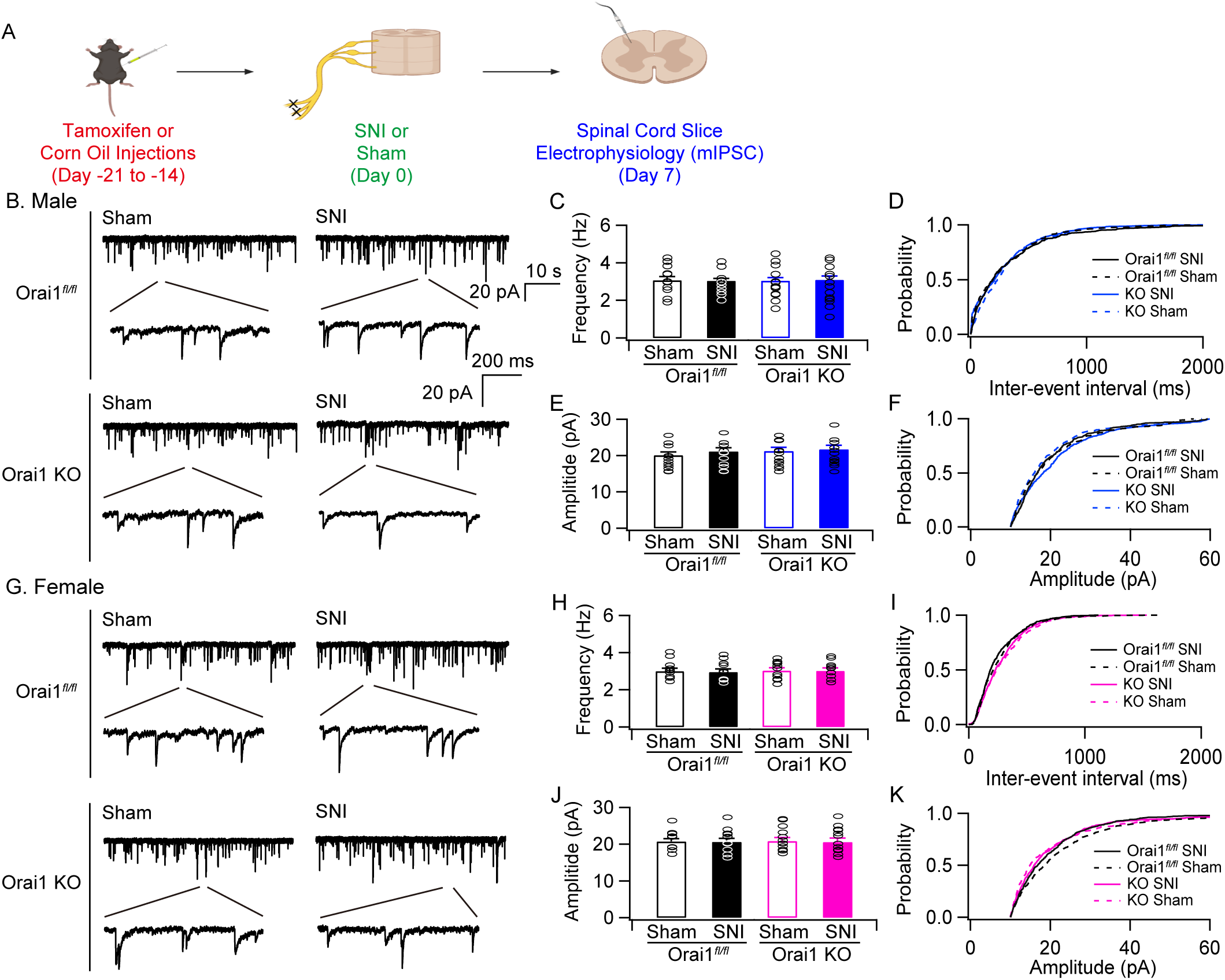
SNI does not affect the frequency and amplitude of miniature IPSCs. mIPSCs were recorded in the whole-cell patch-clamp configuration (-70 mV) from a lamina II neuron in the dorsal horn of the L4 spinal cord in the presence of 1 µM TTX to block action potentials. Excitatory synaptic transmission was blocked by the AMPA and NMDA receptor blockers NBQX and D-APV, respectively. **(B,G)** Example traces of mIPSCs at low and high time resolutions. **(C,E)** The bar graphs show the frequency and amplitude of the mIPCS in the indicated conditions. **(D,F)** Cumulative probability distributions of the inter-event intervals and amplitudes of the mIPSCs in WT and mgOrai1 KO male mice. **(H,J)** Summary of the frequency and amplitude of the mIPCS in female mice in the indicated conditions. **(I,K)** Cumulative probability distributions of the inter-event intervals and amplitudes of the mIPSCs in WT and mgOrai1 KO female mice. n=17 (male); n=12 (female). Statistical analysis was done using one-way ANOVA followed by Tukey test for multiple comparisons.

**Figure S8.**
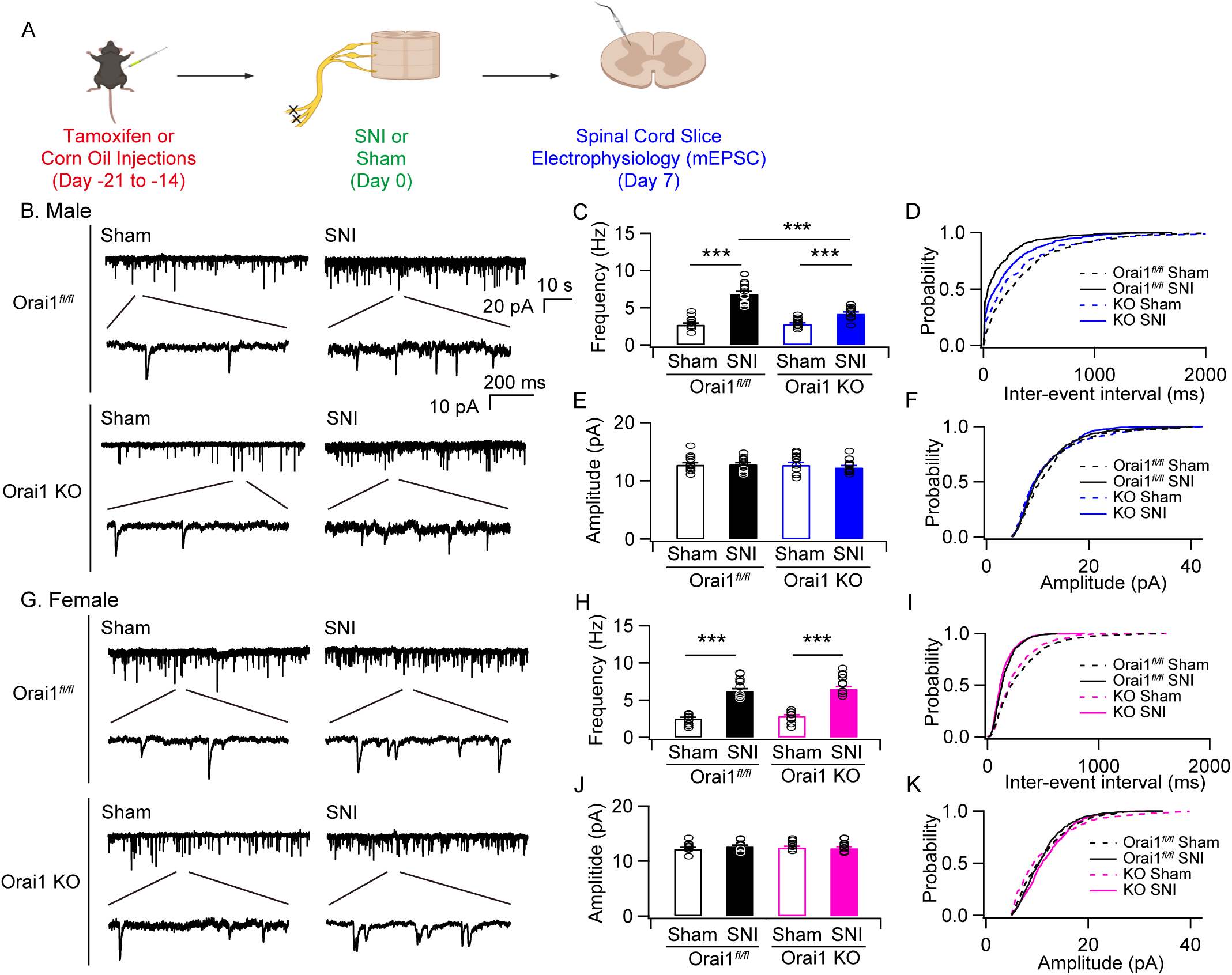
SNI-induced increase in the frequency of miniature EPSCs is mitigated in male mgOrai1 KO mice. **(A)** A schematic of the experiment. WT or mgOrai1 KO mice (7-8 weeks old) were subjected to SNI or sham surgery. 7 days following SNI, animals were euthanized and the spinal cords isolated for slice electrophysiology. **(B,G)** mEPSCs recorded in the whole-cell patch-clamp configuration (-70 mV) from a lamina II neurons in the dorsal horn of the L4 spinal cord. TTX (1 µM) was perfused on the slices in all solutions to block action potentials. The traces show examples of mEPSCs at low and high time resolutions. **(C,E)** Bar graphs summarizing the frequency and amplitude of mEPCS in the indicated conditions. SNI causes increases in the frequency of mEPSCs which is occluded in male mgOrai1 KO mice. There is no change in the amplitude of the events. **(D,F)** Cumulative probability distributions of the inter-event intervals and amplitudes of the mEPSCs in WT and mgOrai1 KO mice. The inter-event interval is shifted towards lower intervals consistent with increases in the frequency of mEPSCs in mice subjected to SNI. **(H,J)** Summary of the change in frequency and amplitude of mEPSCs in female mice. In female mgOrai1 KO mice, the frequency of mEPSC is unaffected and comparable to SNI-administered WT mice. **(I-K)** Cumulative probability distributions of the inter-event intervals and amplitudes of mEPSCs in WT and mgOrai1 female KO mice. n=13 (male), n=10 (female). ***: p<0.001 by ANOVA followed by Tukey test for comparison between multiple groups.

**Figure S9.**
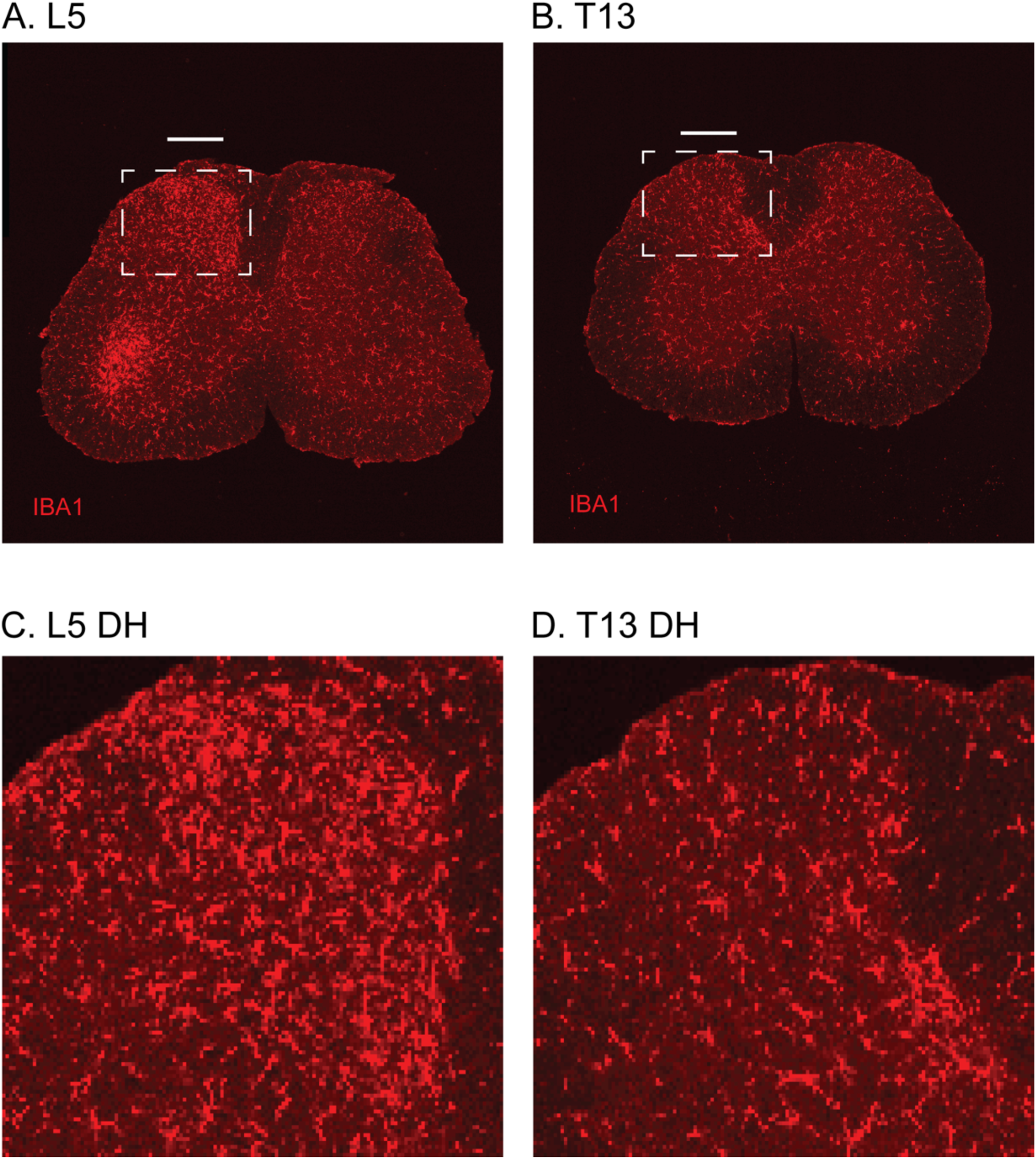
IBA1 Staining in the Lumbar and Thoracic Spinal Cord after SNI. **(A)** Image of the L5 spinal cord from a WT mouse after SNI showing an increase in IBA1 staining in the spinal dorsal horn on the ipsilateral side of injury. **(B)** Image of the thoracic spinal cord at T13 from the same mouse showing IBA1 staining that is evenly distributed throughout the spinal cord. **(C)** Zoomed-in of the ipsilateral side of the spinal dorsal horn after SNI. It shows a clear increase in IBA1 staining after SNI covering the dorsal horn. **(D)** A zoomed in image of the T13 spinal cord area showing IBA1 staining in the ipsilateral dorsal horn.

**Figure S10.**
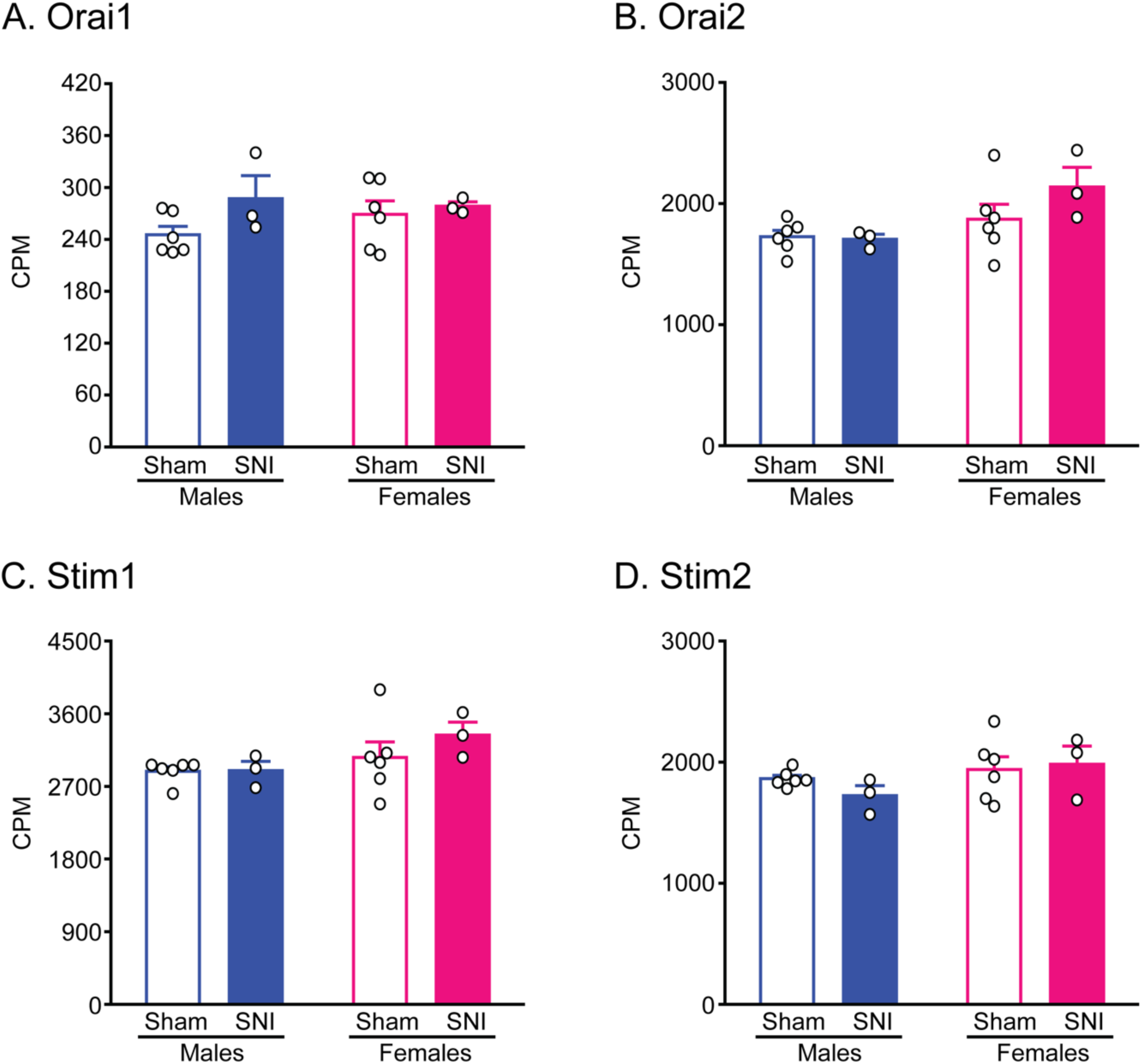
The expression levels of SOCE genes are unchanged after SNI. (A-D) Bar graphs of the normalized counts-per-million (CPM) reads from each sample (L4-L5 spinal cord from one mouse). SNI samples were collected from the ipsilateral dorsal horn of WT mice after SNI. The samples in the sham group contain both ipsilateral and contralateral dorsal horn expression data. After two-way ANOVA analysis of CPM, there is no significant difference in any of the genes between sham and SNI. Additionally, there is no difference between male and female WT mice after SNI. (A) Orai1 (B) Orai2 (C) STIM1 (D) STIM2.

**Figure S11.**
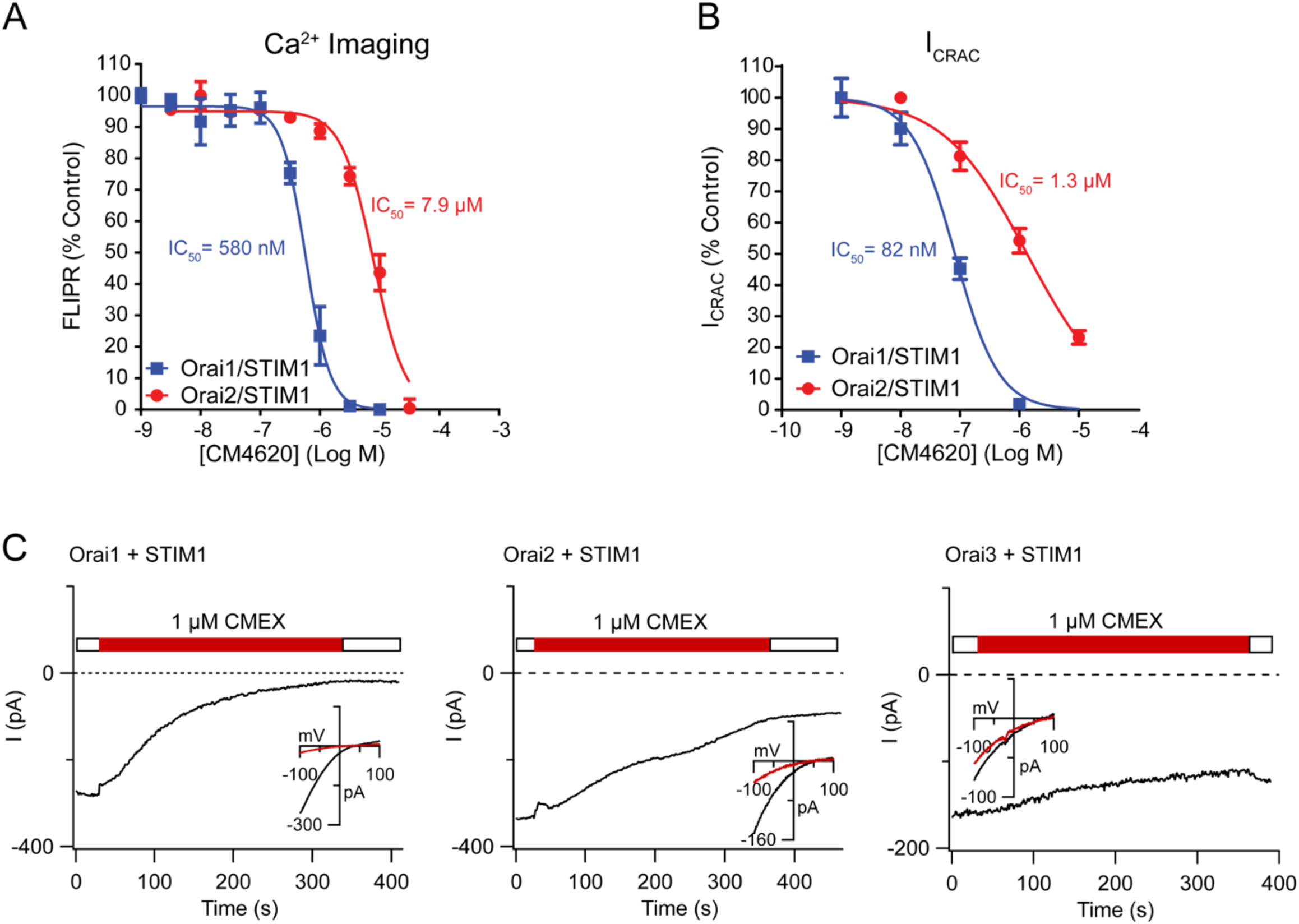
Dose dependence of CM4620 inhibition of Orai1-mediated SOCE and drug specificity. **(A, B)**. Dose-response of CM4620 inhibition of SOCE and CRAC currents measured in HEK293 cells expressing either Orai1 or Orai2 together with STIM1. **(C).** Inhibition of ICRAC by 1 µM CM4620 in HEK293 cells expressing either Orai1, Orai2, or Orai3 together with mCherry-STIM1. The insets show the current-voltage relationship of the Orai current before and after CM4620 administration.

**Figure S12.**
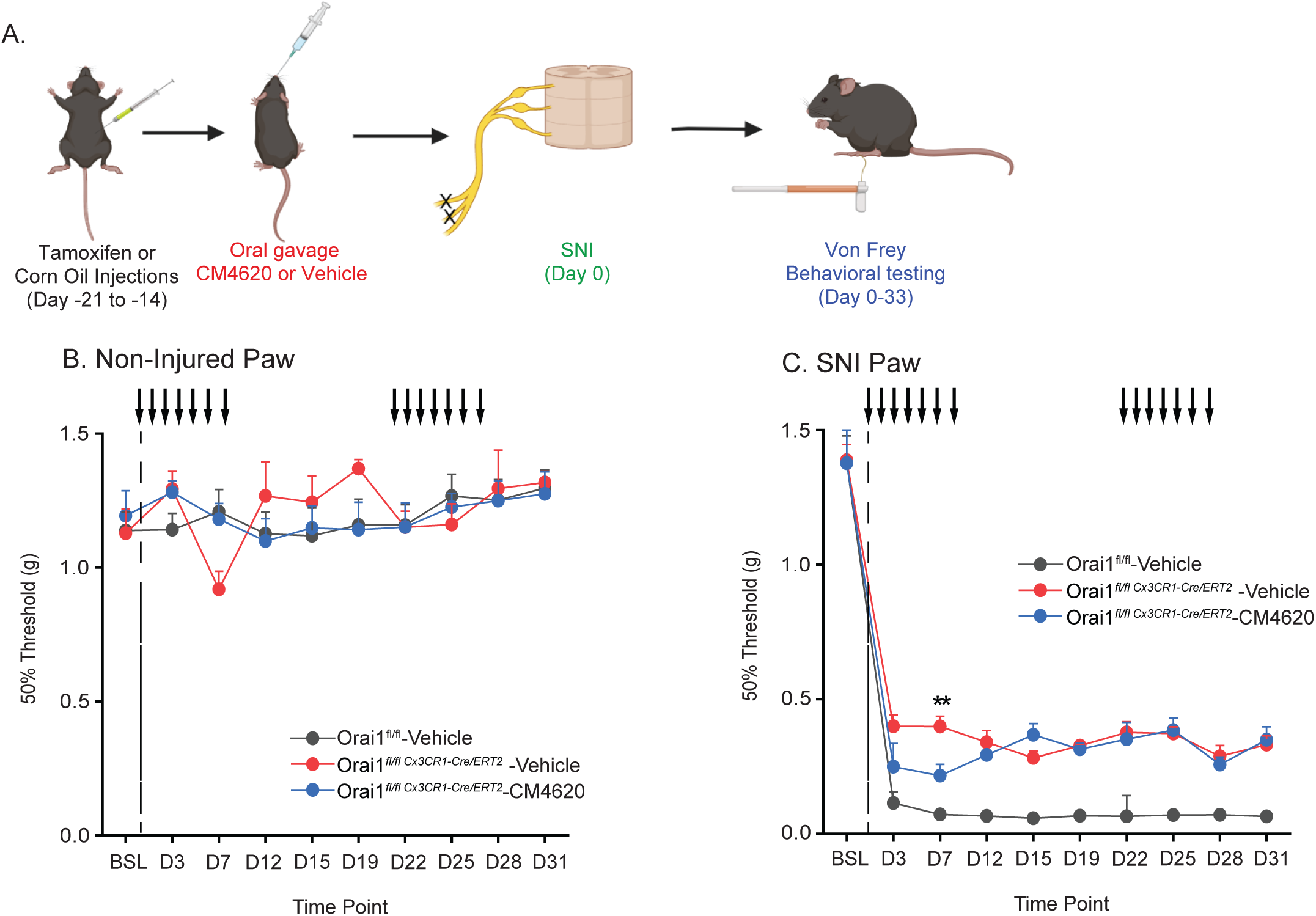
Analysis of pain hypersensitivity in male Orai1*^fl/fl^ ^CX3CR1-Cre/ERT2^* mice administered with CM4620. Mechanical sensitivity was measured by von Frey thresholds in Orai1*^fl/fl^ ^CX3CR1-Cre/ERT2^* and Orai1*^fl/fl^* mice injected with tamoxifen. The dotted line indicates that day when SNI was performed. CM4620 (10 mg/kg) was delivered by oral gavage for 7 days as indicated by the arrows. von Frey thresholds were determined at periodic intervals for 31 days after SNI. CM4620 was stopped after day 8 for 14 days and resumed on day 22. n=7 mice, Orai1*^fl/fl^*(3 males, 4 females); n=6 mice Orai1*^fl/flCX3CR1-Cre^* (all male); n=6 mice, Orai1*^fl/flCX3CR1-Cre^* + CM4620 (all male). **: p < 0.01 by two-way ANOVA followed by Tukey test for comparison between multiple groups at each time point.

## REFERENCES

1. Q. Li, B. A. Barres, Microglia and macrophages in brain homeostasis and disease. Nat Rev Immunol 18, 225–242 (2018).

2. W. J. Streit, S. A. Walter, N. A. Pennell, Reactive microgliosis. Progress in neurobiology 57, 563–581 (1999).

3. M. W. Salter, B. Stevens, Microglia emerge as central players in brain disease. Nat Med 23, 1018–1027 (2017).

4. K. Inoue, M. Tsuda, Microglia in neuropathic pain: cellular and molecular mechanisms and therapeutic potential. Nat Rev Neurosci 19, 138–152 (2018).

5. M. Tsuda, S. Beggs, M. W. Salter, K. Inoue, Microglia and intractable chronic pain. Glia 61, 55–61 (2013).

6. S. Hickman, S. Izzy, P. Sen, L. Morsett, J. El Khoury, Microglia in neurodegeneration. Nat Neurosci 21, 1359–1369 (2018).

7. T. Trang, S. Beggs, M. W. Salter, ATP receptors gate microglia signaling in neuropathic pain. Exp Neurol 234, 354–361 (2012).

8. T. Masuda, Y. Ozono, S. Mikuriya, Y. Kohro, H. Tozaki-Saitoh, K. Iwatsuki, H. Uneyama, R. Ichikawa, M. W. Salter, M. Tsuda, K. Inoue, Dorsal horn neurons release extracellular ATP in a VNUT-dependent manner that underlies neuropathic pain. Nat Commun 7, 12529 (2016).

9. N. Gu, U. B. Eyo, M. Murugan, J. Peng, S. Matta, H. Dong, L. J. Wu, Microglial P2Y12 receptors regulate microglial activation and surveillance during neuropathic pain. Brain Behav Immun 55, 82–92 (2016).

10. H. Tozaki-Saitoh, M. Tsuda, H. Miyata, K. Ueda, S. Kohsaka, K. Inoue, P2Y12 receptors in spinal microglia are required for neuropathic pain after peripheral nerve injury. J Neurosci 28, 4949–4956 (2008).

11. X. Zhang, G. Li, P2Y receptors in neuropathic pain. Pharmacol Biochem Behav 186, 172788 (2019).

12. K. Kohno, M. Tsuda, Role of microglia and P2X4 receptors in chronic pain. Pain Rep 6, e864 (2021).

13. X. Yan, H. R. Weng, Endogenous interleukin-1beta in neuropathic rats enhances glutamate release from the primary afferents in the spinal dorsal horn through coupling with presynaptic N-methyl-D-aspartic acid receptors. J Biol Chem 288, 30544–30557 (2013).

14. G. Chen, Y. Q. Zhang, Y. J. Qadri, C. N. Serhan, R. R. Ji, Microglia in Pain: Detrimental and Protective Roles in Pathogenesis and Resolution of Pain. Neuron 100, 1292–1311 (2018).

15. Y. Kawasaki, L. Zhang, J. K. Cheng, R. R. Ji, Cytokine mechanisms of central sensitization: distinct and overlapping role of interleukin-1beta, interleukin-6, and tumor necrosis factor-alpha in regulating synaptic and neuronal activity in the superficial spinal cord. J Neurosci 28, 5189–5194 (2008).

16. R. R. Ji, A. Chamessian, Y. Q. Zhang, Pain regulation by non-neuronal cells and inflammation. Science 354, 572–577 (2016).

17. R. R. Ji, A. Nackley, Y. Huh, N. Terrando, W. Maixner, Neuroinflammation and Central Sensitization in Chronic and Widespread Pain. Anesthesiology 129, 343–366 (2018).

18. Y. Q. Zhou, Z. Liu, Z. H. Liu, S. P. Chen, M. Li, A. Shahveranov, D. W. Ye, Y. K. Tian, Interleukin-6: an emerging regulator of pathological pain. J Neuroinflammation 13, 141 (2016).

19. E. Gabay, G. Wolf, Y. Shavit, R. Yirmiya, M. Tal, Chronic blockade of interleukin-1 (IL-1) prevents and attenuates neuropathic pain behavior and spontaneous ectopic neuronal activity following nerve injury. Eur J Pain 15, 242–248 (2011).

20. L. Leung, C. M. Cahill, TNF-alpha and neuropathic pain--a review. J Neuroinflammation 7, 27 (2010).

21. K. Serizawa, H. Tomizawa-Shinohara, H. Yasuno, K. Yogo, Y. Matsumoto, Anti-IL-6 Receptor Antibody Inhibits Spontaneous Pain at the Pre-onset of Experimental Autoimmune Encephalomyelitis in Mice. Frontiers in neurology 10, 341 (2019).

22. V. Raghavendra, F. Tanga, J. A. DeLeo, Inhibition of microglial activation attenuates the development but not existing hypersensitivity in a rat model of neuropathy. J Pharmacol Exp Ther 306, 624–630 (2003).

23. R. E. Sorge, J. C. Mapplebeck, S. Rosen, S. Beggs, S. Taves, J. K. Alexander, L. J. Martin, J. S. Austin, S. G. Sotocinal, D. Chen, M. Yang, X. Q. Shi, H. Huang, N. J. Pillon, P. J. Bilan, Y. Tu, A. Klip, R. R. Ji, J. Zhang, M. W. Salter, J. S. Mogil, Different immune cells mediate mechanical pain hypersensitivity in male and female mice. Nat Neurosci 18, 1081–1083 (2015).

24. A. Ledeboer, E. M. Sloane, E. D. Milligan, M. G. Frank, J. H. Mahony, S. F. Maier, L. R. Watkins, Minocycline attenuates mechanical allodynia and proinflammatory cytokine expression in rat models of pain facilitation. Pain 115, 71–83 (2005).

25. J. C. Mapplebeck, S. Beggs, M. W. Salter, Sex differences in pain: a tale of two immune cells. Pain 157 **Suppl 1**, S2–6 (2016).

26. S. Taves, T. Berta, D. L. Liu, S. Gan, G. Chen, Y. H. Kim, T. Van de Ven, S. Laufer, R. R. Ji, Spinal inhibition of p38 MAP kinase reduces inflammatory and neuropathic pain in male but not female mice: Sex-dependent microglial signaling in the spinal cord. Brain Behav Immun 55, 70–81 (2016).

27. J. C. S. Mapplebeck, R. Dalgarno, Y. Tu, O. Moriarty, S. Beggs, C. H. T. Kwok, K. Halievski, S. Assi, J. S. Mogil, T. Trang, M. W. Salter, Microglial P2X4R-evoked pain hypersensitivity is sexually dimorphic in rats. Pain 159, 1752–1763 (2018).

28. S. Rosen, B. Ham, J. S. Mogil, Sex differences in neuroimmunity and pain. J Neurosci Res 95, 500–508 (2017).

29. K. Halievski, S. Ghazisaeidi, M. W. Salter, Sex-Dependent Mechanisms of Chronic Pain: A Focus on Microglia and P2X4R. J Pharmacol Exp Ther 375, 202–209 (2020).

30. C. J. Woolf, What is this thing called pain? J Clin Invest 120, 3742–3744 (2010).

31. L. Colloca, T. Ludman, D. Bouhassira, R. Baron, A. H. Dickenson, D. Yarnitsky, R. Freeman, A. Truini, N. Attal, N. B. Finnerup, C. Eccleston, E. Kalso, D. L. Bennett, R. H. Dworkin, S. N. Raja, Neuropathic pain. Nature reviews. Disease primers 3, 17002 (2017).

32. M. N. Baliki, A. V. Apkarian, Nociception, Pain, Negative Moods, and Behavior Selection. Neuron 87, 474–491 (2015).

33. K. Farber, H. Kettenmann, Functional role of calcium signals for microglial function. Glia 54, 656–665 (2006).

34. T. Moller, Calcium signaling in microglial cells. Glia 40, 184–194 (2002).

35. S. Zumerle, B. Cali, F. Munari, R. Angioni, F. Di Virgilio, B. Molon, A. Viola, Intercellular Calcium Signaling Induced by ATP Potentiates Macrophage Phagocytosis. Cell reports 27, 1–10 e14 (2019).

36. A. Mizuma, J. Y. Kim, R. Kacimi, K. Stauderman, M. Dunn, S. Hebbar, M. A. Yenari, Microglial Calcium Release-Activated Calcium Channel Inhibition Improves Outcome from Experimental Traumatic Brain Injury and Microglia-Induced Neuronal Death. Journal of Neurotrauma 36, 996–1007 (2019).

37. M. Prakriya, R. S. Lewis, Store-Operated Calcium Channels. Physiol Rev 95, 1383–1436 (2015).

38. P. S. Yeung, M. Yamashita, M. Prakriya, Molecular basis of allosteric Orai1 channel activation by STIM1. J Physiol 598, 1707–1723 (2020).

39. M. Michaelis, B. Nieswandt, D. Stegner, J. Eilers, R. Kraft, STIM1, STIM2, and Orai1 regulate store-operated calcium entry and purinergic activation of microglia. Glia 63, 652–663 (2015).

40. D. F. Gilbert, M. J. Stebbing, K. Kuenzel, R. M. Murphy, E. Zacharewicz, A. Buttgereit, L. Stokes, D. J. Adams, O. Friedrich, Store-Operated Ca^2+^ Entry (SOCE) and Purinergic Receptor-Mediated Ca^2+^ Homeostasis in Murine bv2 Microglia Cells: Early Cellular Responses to ATP-Mediated Microglia Activation. Frontiers Molecular Neuroscience 9, 111 (2016).

41. H. M. Lim, H. Woon, J. W. Han, Y. Baba, T. Kurosaki, M. G. Lee, J. Y. Kim, UDP-Induced Phagocytosis and ATP-Stimulated Chemotactic Migration Are Impaired in STIM1(-/-) Microglia In Vitro and In Vivo. Mediators Inflamm 2017, 8158514 (2017).

42. A. B. Toth, K. Hori, M. M. Novakovic, N. G. Bernstein, L. Lambot, M. Prakriya, CRAC channels regulate astrocyte Ca^2+^ signaling and gliotransmitter release to modulate hippocampal GABAergic transmission. Sci Signal 12 eaaw5450, 1–15 (2019).

43. M. M. Maneshi, A. B. Toth, T. Ishii, K. Hori, S. Tsujikawa, A. K. Shum, N. Shrestha, M. Yamashita, R. J. Miller, J. Radulovic, G. T. Swanson, M. Prakriya, Orai1 Channels Are Essential for Amplification of Glutamate-Evoked Ca^2+^ Signals in Dendritic Spines to Regulate Working and Associative Memory. Cell reports 33, 108464 (2020).

44. Y. Dou, J. Xia, R. Gao, X. Gao, F. M. Munoz, D. Wei, Y. Tian, J. E. Barrett, S. Ajit, O. Meucci, J. W. Putney, Jr., Y. Dai, H. Hu, Orai1 Plays a Crucial Role in Central Sensitization by Modulating Neuronal Excitability. J Neurosci 38, 887–900 (2018).

45. X. F. Zhao, M. M. Alam, Y. Liao, T. Huang, R. Mathur, X. Zhu, Y. Huang, Targeting Microglia Using Cx3cr1-Cre Lines: Revisiting the Specificity. eNeuro 6, ENEURO.0114-19.2019. doi: 10.1523/ENEURO.0114-19.2019 (2019).

46. A. Somasundaram, A. K. Shum, H. J. McBride, J. A. Kessler, S. Feske, R. J. Miller, M. Prakriya, Store-operated CRAC channels regulate gene expression and proliferation in neural progenitor cells. J Neurosci 34, 9107–9123 (2014).

47. R. T. Waldron, Y. Chen, H. Pham, A. Go, H. Y. Su, C. Hu, L. Wen, S. Z. Husain, C. A. Sugar, J. Roos, S. Ramos, A. Lugea, M. Dunn, K. Stauderman, S. J. Pandol, The Orai Ca^2+^ channel inhibitor CM4620 targets both parenchymal and immune cells to reduce inflammation in experimental acute pancreatitis. J Physiol 597, 3085–3105 (2019).

48. J. Miller, C. Bruen, M. Schnaus, J. Zhang, S. Ali, A. Lind, Z. Stoecker, K. Stauderman, S. Hebbar, Auxora versus standard of care for the treatment of severe or critical COVID-19 pneumonia: results from a randomized controlled trial. Crit Care 24, 502 (2020).

49. S. M. Emrich, R. E. Yoast, M. Trebak, Physiological Functions of CRAC Channels. Annu Rev Physiol, (2021).

50. T. S. Kountz, A. Jairaman, C. D. Kountz, K. A. Stauderman, R. P. Schleimer, M. Prakriya, Differential Regulation of ATP- and UTP-Evoked Prostaglandin E2 and IL-6 Production from Human Airway Epithelial Cells. J Immunol 207, 1275–1287 (2021).

51. R. E. Sorge, M. L. LaCroix-Fralish, A. H. Tuttle, S. G. Sotocinal, J. S. Austin, J. Ritchie, M. L. Chanda, A. C. Graham, L. Topham, S. Beggs, M. W. Salter, J. S. Mogil, Spinal cord Toll-like receptor 4 mediates inflammatory and neuropathic hypersensitivity in male but not female mice. J Neurosci 31, 15450–15454 (2011).

52. T. Liu, Q. Han, G. Chen, Y. Huang, L. X. Zhao, T. Berta, Y. J. Gao, R. R. Ji, Toll-like receptor 4 contributes to chronic itch, alloknesis, and spinal astrocyte activation in male mice. Pain 157, 806–817 (2016).

53. F. Y. Tanga, N. Nutile-McMenemy, J. A. DeLeo, The CNS role of Toll-like receptor 4 in innate neuroimmunity and painful neuropathy. Proc Natl Acad Sci U S A 102, 5856–5861 (2005).

54. M. J. Lacagnina, L. R. Watkins, P. M. Grace, Toll-like receptors and their role in persistent pain. Pharmacology & therapeutics 184, 145–158 (2018).

55. A. Beck, R. Penner, A. Fleig, Lipopolysaccharide-induced down-regulation of Ca^2+^ release-activated Ca2+ currents (I CRAC) but not Ca^2+^-activated TRPM4-like currents (I CAN) in cultured mouse microglial cells. J Physiol 586, 427–439 (2008).

56. P. G. Hogan, L. Chen, J. Nardone, A. Rao, Transcriptional regulation by calcium, calcineurin, and NFAT. Genes Dev 17, 2205–2232 (2003).

57. J. K. Olson, S. D. Miller, Microglia initiate central nervous system innate and adaptive immune responses through multiple TLRs. J Immunol 173, 3916–3924 (2004).

58. T. Wang, L. Qin, B. Liu, Y. Liu, B. Wilson, T. E. Eling, R. Langenbach, S. Taniura, J. S. Hong, Role of reactive oxygen species in LPS-induced production of prostaglandin E2 in microglia. J Neurochem 88, 939–947 (2004).

59. D. Gruber-Schoffnegger, R. Drdla-Schutting, C. Honigsperger, G. Wunderbaldinger, M. Gassner, J. Sandkuhler, Induction of thermal hyperalgesia and synaptic long-term potentiation in the spinal cord lamina I by TNF-alpha and IL-1beta is mediated by glial cells. J Neurosci 33, 6540–6551 (2013).

60. A. V. Apkarian, A. A. Mutso, M. V. Centeno, L. Kan, M. Wu, M. Levinstein, G. Banisadr, K. T. Gobeske, R. J. Miller, J. Radulovic, R. Hen, J. A. Kessler, Role of adult hippocampal neurogenesis in persistent pain. Pain 157, 418–428 (2016).

61. I. Decosterd, C. J. Woolf, Spared nerve injury: an animal model of persistent peripheral neuropathic pain. Pain 87, 149–158 (2000).

62. C. N. Parkhurst, G. Yang, I. Ninan, J. N. Savas, J. R. Yates, 3rd, J. J. Lafaille, B. L. Hempstead, D. R. Littman, W. B. Gan, Microglia promote learning-dependent synapse formation through brain-derived neurotrophic factor. Cell 155, 1596–1609 (2013).

63. R. van Furth, Z. A. Cohn, The origin and kinetics of mononuclear phagocytes. J Exp Med 128, 415–435 (1968).

64. D. K. Fogg, C. Sibon, C. Miled, S. Jung, P. Aucouturier, D. R. Littman, A. Cumano, F. Geissmann, A clonogenic bone marrow progenitor specific for macrophages and dendritic cells. Science 311, 83–87 (2006).

65. B. Ajami, J. L. Bennett, C. Krieger, W. Tetzlaff, F. M. Rossi, Local self-renewal can sustain CNS microglia maintenance and function throughout adult life. Nat Neurosci 10, 1538–1543 (2007).

66. P. Reu, A. Khosravi, S. Bernard, J. E. Mold, M. Salehpour, K. Alkass, S. Perl, J. Tisdale, G. Possnert, H. Druid, J. Frisen, The Lifespan and Turnover of Microglia in the Human Brain. Cell reports 20, 779–784 (2017).

67. W. Renthal, I. Tochitsky, L. Yang, Y. C. Cheng, E. Li, R. Kawaguchi, D. H. Geschwind, C. J. Woolf, Transcriptional Reprogramming of Distinct Peripheral Sensory Neuron Subtypes after Axonal Injury. Neuron 108, 128–144 e129 (2020).

68. T. Murakami, T. Kanchiku, H. Suzuki, Y. Imajo, Y. Yoshida, H. Nomura, D. Cui, T. Ishikawa, E. Ikeda, T. Taguchi, Anti-interleukin-6 receptor antibody reduces neuropathic pain following spinal cord injury in mice. Exp Ther Med 6, 1194–1198 (2013).

69. Y. Nakajima, K. Osuka, Y. Seki, R. C. Gupta, M. Hara, M. Takayasu, T. Wakabayashi, Taurine reduces inflammatory responses after spinal cord injury. Journal of Neurotrauma 27, 403–410 (2010).

70. Y. Liu, L. J. Zhou, J. Wang, D. Li, W. J. Ren, J. Peng, X. Wei, T. Xu, W. J. Xin, R. P. Pang, Y. Y. Li, Z. H. Qin, M. Murugan, M. P. Mattson, L. J. Wu, X. G. Liu, TNF-alpha Differentially Regulates Synaptic Plasticity in the Hippocampus and Spinal Cord by Microglia-Dependent Mechanisms after Peripheral Nerve Injury. J Neurosci 37, 871–881 (2017).

71. J. Sandkuhler, D. Gruber-Schoffnegger, Hyperalgesia by synaptic long-term potentiation (LTP): an update. Curr Opin Pharmacol 12, 18–27 (2012).

72. L. Zhang, T. Berta, Z. Z. Xu, T. Liu, J. Y. Park, R. R. Ji, TNF-alpha contributes to spinal cord synaptic plasticity and inflammatory pain: distinct role of TNF receptor subtypes 1 and 2. Pain 152, 419–427 (2011).

73. A. K. Clark, D. Gruber-Schoffnegger, R. Drdla-Schutting, K. J. Gerhold, M. Malcangio, J. Sandkuhler, Selective activation of microglia facilitates synaptic strength. J Neurosci 35, 4552–4570 (2015).

74. D. Uta, G. Kato, A. Doi, T. Andoh, T. Kume, M. Yoshimura, K. Koga, Animal models of chronic pain increase spontaneous glutamatergic transmission in adult rat spinal dorsal horn in vitro and in vivo. Biochem Biophys Res Commun 512, 352–359 (2019).

75. Y. Hayano, K. Takasu, Y. Koyama, M. Yamada, K. Ogawa, K. Minami, T. Asaki, K. Kitada, S. Kuwabara, T. Yamashita, Dorsal horn interneuron-derived Netrin-4 contributes to spinal sensitization in chronic pain via Unc5B. J Exp Med 213, 2949–2966 (2016).

76. X. Yan, E. Jiang, M. Gao, H. R. Weng, Endogenous activation of presynaptic NMDA receptors enhances glutamate release from the primary afferents in the spinal dorsal horn in a rat model of neuropathic pain. J Physiol 591, 2001–2019 (2013).

77. K. Hori, S. Tsujikawa, M. M. Novakovic, M. Yamashita, M. Prakriya, Regulation of chemoconvulsant-induced seizures by store-operated Orai1 channels. J Physiol 598, 5391–5409 (2020).

78. T. Yu, X. Zhang, H. Shi, J. Tian, L. Sun, X. Hu, W. Cui, D. Du, P2Y12 regulates microglia activation and excitatory synaptic transmission in spinal lamina II neurons during neuropathic pain in rodents. Cell death & disease 10, 165 (2019).

79. M. Deng, S. R. Chen, H. L. Pan, Presynaptic NMDA receptors control nociceptive transmission at the spinal cord level in neuropathic pain. Cellular and Molecular Life Sciences : CMLS 76, 1889–1899 (2019).

80. S. Ghazisaeidi, M. M. Muley, Y. Tu, M. Kolahdouzan, A. S. Sengar, A. K. Ramani, M. Brudno, M. W. Salter, Conserved transcriptional programming across sex and species after peripheral nerve injury predicts treatments for neuropathic pain. bioRxiv, 2022.2005.2030.494054 (2022).

81. R. Gao, X. Gao, J. Xia, Y. Tian, J. E. Barrett, Y. Dai, H. Hu, Potent analgesic effects of a store-operated calcium channel inhibitor. Pain 154, 2034–2044 (2013).

82. R. Takezawa, H. Cheng, A. Beck, J. Ishikawa, P. Launay, H. Kubota, J. P. Kinet, A. Fleig, T. Yamada, R. Penner, A pyrazole derivative potently inhibits lymphocyte Ca^2+^ influx and cytokine production by facilitating transient receptor potential melastatin 4 channel activity. Mol Pharmacol 69, 1413–1420 (2006).

83. A. Meizoso-Huesca, B. S. Launikonis, The Orai1 inhibitor BTP2 has multiple effects on Ca2+ handling in skeletal muscle. J Gen Physiol 153, (2021).

84. M. Vaeth, S. Kahlfuss, S. Feske, CRAC Channels and Calcium Signaling in T Cell-Mediated Immunity. Trends Immunol 41, 878–901 (2020).

85. S. Feske, CRAC channels and disease - From human CRAC channelopathies and animal models to novel drugs. Cell Calcium 80, 112–116 (2019).

86. D. K. Heo, H. M. Lim, J. H. Nam, M. G. Lee, J. Y. Kim, Regulation of phagocytosis and cytokine secretion by store-operated calcium entry in primary isolated murine microglia. Cell Signal 27, 177–186 (2015).

87. K. Inoue, The function of microglia through purinergic receptors: Neuropathic pain and cytokine release. Pharmacology & Therapeutics 109, 210–226 (2006).

88. S. T. T. Schetters, D. Gomez-Nicola, J. J. Garcia-Vallejo, Y. Van Kooyk, Neuroinflammation: Microglia and T Cells Get Ready to Tango. Frontiers in Immunology 8, 1905 (2017).

89. T. Zoller, A. Schneider, C. Kleimeyer, T. Masuda, P. S. Potru, D. Pfeifer, T. Blank, M. Prinz, B. Spittau, Silencing of TGFbeta signalling in microglia results in impaired homeostasis. Nat Commun 9, 4011 (2018).

90. J. K. Lee, M. G. Tansey, Microglia isolation from adult mouse brain. Methods in molecular biology 1041, 17–23 (2013).

91. S. R. Chaplan, F. W. Bach, J. W. Pogrel, J. M. Chung, T. L. Yaksh, Quantitative assessment of tactile allodynia in the rat paw. J Neurosci Methods 53, 55–63 (1994).

